# Cancer immunotherapy by NC410, a LAIR-2 Fc protein blocking LAIR-collagen interaction

**DOI:** 10.1101/2020.10.21.349480

**Authors:** M. Inês Pascoal Ramos, Linjie Tian, Emma J. de Ruiter, Chang Song, Ana Paucarmayta, Akashdip Singh, Eline Elshof, Saskia V. Vijver, Jahangheer Shaik, Jason Bosiacki, Zachary Cusumano, Linda Liu, Sol Langermann, Stefan Willems, Dallas Flies, Linde Meyaard

**Affiliations:** Center for Translational Immunology, University Medical Center Utrecht, Utrecht University, Utrecht, The Netherlands; Oncode Institute, Utrecht, The Netherlands; NextCure, Beltsville, MD, USA; Department of Pathology, University Medical Center Utrecht, Utrecht University, Utrecht, The Netherlands

## Abstract

Collagens are a primary component of the extracellular matrix and are functional ligands for the inhibitory immune receptor leukocyte associated immunoglobulin-like receptor-1 (LAIR-1). Leukocyte associated immunoglobulin-like receptor-2 (LAIR-2) is a secreted protein that can act as a decoy receptor by binding collagen with higher affinity than LAIR-1. We propose that collagens promote immune evasion by interacting with LAIR-1 and that LAIR-2 could release LAIR-1 mediated immune suppression. Analysis of public datasets shows high LAIR-2 expression being associated with a favorable outcome in certain tumors. We designed a dimeric LAIR-2 with a functional IgG1 Fc tail, NC410, and showed that NC410 reduces tumor growth and increases T cell expansion and effector function in humanized tumor models. Immunohistochemical analysis of human tumors shows that NC410 binds to collagen-rich areas where LAIR-1^+^ immune cells are localized. Our findings show that NC410 might be a powerful new strategy for cancer immunotherapy for immune-excluded tumors.

## Introduction

The introduction of immune checkpoint blockade therapies in the clinic has increased cancer treatment options for a wide range of tumors leading to unprecedented and long-lasting clinical responses. However, not all patients show the same degree of response and not all tumors respond to these therapies (1, 2). Thus, identifying novel checkpoints and developing ideal combinations of immunotherapies is essential to optimize and enhance the efficacy of treatment, achieving durable anti-cancer effects with reduced side effects.

The extracellular matrix (ECM) is a major structural component in all tissues. It comprises a non-cellular meshwork of proteins, glycoproteins, proteoglycans and polysaccharides with collagens as the most abundant protein. Ongoing ECM remodeling ensures tissue integrity and function, with collagens being synthesized and degraded in a highly regulated manner (3). At least 28 different collagens comprised of at least 43 genes have been identified from which 4 have transmembrane domains allowing expression on the cell surface (4). The ECM functions not only as a scaffold for tissue organization but also provides critical biochemical and biomechanical cues that instruct cell growth, survival, differentiation and migration and regulate vascular development and immune function (5).

Several epithelial cancers including breast, pancreatic, colorectal, ovarian, and lung cancer are characterized by a dense ECM where high collagen content correlates with poor prognosis (6). Indeed, ECM or “Matrisome” signatures associated with cancer type and stage of disease have been described (7–10). Cancer associated fibroblasts (CAFs) (11, 12), macrophages and tumor cells themselves (7, 13, 14) all contribute to increased collagen production during cancer progression.

The capacity of tumors to induce remodeling of collagens in the tumor microenvironment (TME) was primarily thought to create a suitable microenvironment for tumor cell growth. We now consider abnormal collagen production, composition and organization in the ECM-TME a cause of immune dysfunction, conveying a chronic wound-healing response instead of anti-tumor immune responses necessary for immune surveillance and the eradication of the tumor (15, 16). The abnormal ECM also builds physical barriers to exclude immune cells and therapeutic agents from access to tumor cells (17). Furthermore, tumor-associated collagen can interact with the inhibitory collagen receptor leukocyte associated immunoglobulin-like receptor-1 (LAIR-1)(18). Other collagen-domain containing LAIR-1 ligands have also been reported to be present and enriched in the TME (19). LAIR-1 is an immune checkpoint broadly expressed on the cell surface of immune cells (20) that binds to collagen (21) and molecules with collagen like domains (22, 23). LAIR-1^+^ cells strongly adhere to collagen (21). Upon triggering, LAIR-1 inhibits NK cell (24), T cell (25–27), B cell (28, 29), monocyte (30) and DC function (31, 32). Thus, besides formation of a tumor niche, tumor-associated collagens can function to promote immune evasion through its interaction with LAIR-1.

We sought to develop an immunomedicine that targets the link between tumor ECM abnormalities and T cell dysfunction to resolve immune suppression and improve tumor clearance. We utilized leukocyte associated immunoglobulin-like receptor-2 (LAIR-2), a soluble collagen binding protein that can act as a decoy for LAIR-1 and as such is a natural immune checkpoint inhibitor (33, 34). We developed a dimeric LAIR-2 Fc fusion protein, NC410, as a novel immunomedicine to both target tumor ECM and promote T cell activation through blockade of LAIR-1 mediated inhibition.

## Results

### Collagen and LAIR-1 are overexpressed in tumors and correlate with poor overall survival while overexpression of the decoy receptor LAIR-2 associates with increased survival

Most cancers overexpress a diverse set of collagens in an abnormal fashion (35, 36). To characterize collagen mRNA expression in cancer, we performed a meta-analysis of the TCGA database. 28 collagens comprised of 43 genes were assessed as a group for mRNA expression in cancer and normal tissues across 21 TCGA cancer subtypes. Overexpression of collagens has been associated with poor overall survival in several cancer types, such as lung (37, 38), colorectal (10, 39) and ovarian cancer (40, 41). When we examined the mRNA expression of all collagens genes combined, 13 out of 21 tumors, namely breast invasive carcinoma (BRCA), cholangiocarcinoma (CHOL), colon adenocarcinoma (COAD), esophageal (ESCA), glioblastoma (GBM), head and neck squamous cell carcinoma (HNSC), kidney renal clear cell carcinoma (KIRC), liver hepatocellular carcinoma (LIHC), lung adenocarcinoma (LUAD), lung squamous cell carcinoma (LUSC), rectum adenocarcinoma (READ), stomach adenocarcinoma (STAD) and thyroid carcinoma (THCA) showed increased collagen mRNA expression compared to healthy tissue (Figure 1A top graph). Importantly, this mRNA collagen signature correlated with worse overall survival in 9 out of 21 cancer types analyzed, namely uterine (UCEC), adrenocortical (ACC), low-grade glioma (LGG), sarcoma (SARC), pancreatic (PAAD), esophageal (ESCA), lung (LUAD) and prostate (PRAD) tumors as well as in acute myeloid leukemia (LAML) (Figure 1B). Thus, increased collagen expression and poor clinical outcome associated with overexpression of collagens in multiple cancer types provides a preliminary rationale for targeting collagens as a therapeutic strategy.

**Figure1.**
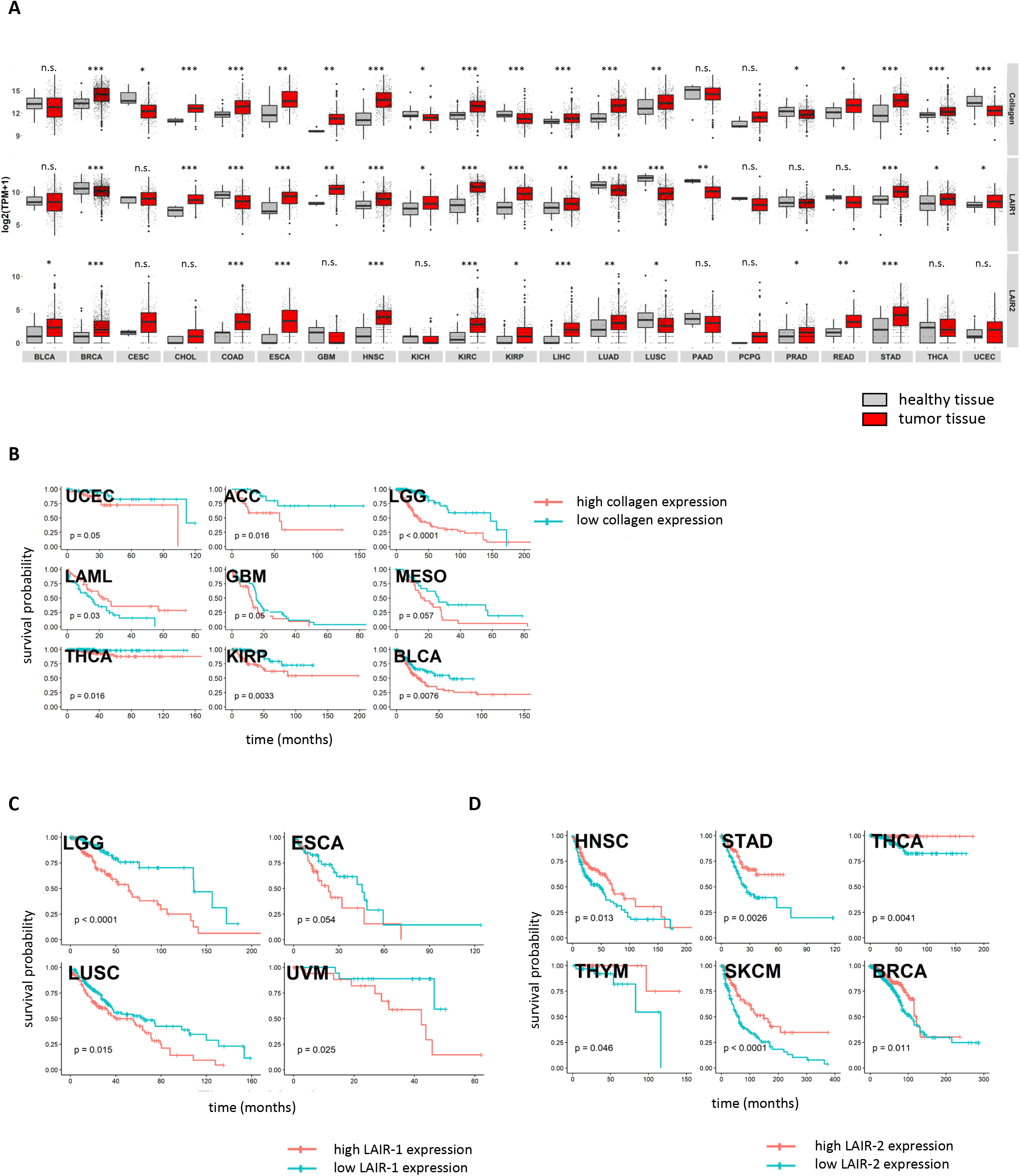
Collagen and LAIR-1 are overexpressed in tumors and correlate with poor overall survival while the decoy receptor LAIR-2 associates with increased survival. A) The expression of 43 collagen genes, LAIR-1 and LAIR-2 in normal (grey) and tumor tissue (red) were queried using TCGA database. B) The log2 transformed collagen expression for each of the 43 collagen proteins in various cancers were divided into four quantiles. The patients in lower quantile (blue) were considered individuals with low expression and those in the upper quantile (red) were considered those with high expression respectively. The estimate of overall survival was determined using Kaplan-Meier method. C) Patients were grouped in low 25% quantile (blue) and high 25% quantile (red) LAIR-1 or D) LAIR-2 mRNA expression for overall survival analysis.

Tumor-expressed collagens can modulate immune cell function through the inhibitory collagen receptor LAIR-1 (18). In the TCGA data, LAIR-1 mRNA expression was significantly upregulated in 11 out of 21 human cancer types when compared to matched healthy tissues or non-tumoral adjacent healthy tissues (Figure 1A, middle graph). By stratifying patients for high and low LAIR-1 mRNA expression we observed that patients with high LAIR-1 mRNA expression had lower survival probability in 4 out of 21 cancers, namely LGG, ESCA, LUAD and LUSC (Figure 1C). We also observed that LAIR-2 mRNA expression, despite being at lower levels than collagen and LAIR-1, was significantly upregulated in 14 out 21 tumors compared to healthy tissue (Figure 1A bottom graph). We posited that increased mRNA expression of LAIR-2, the soluble decoy for the LAIR-1 inhibitory receptor, may be associated with improved overall survival. When patients were stratified by high and low LAIR-2 mRNA expression, high expression of LAIR-2 mRNA was indeed associated with increased overall survival probability in 6 out of 30 cancers analysed, namely HNSC, STAD, thyroid carcinoma (THCA), thymoma (THYM), skin cutaneous melanoma (SKCM) and breast invasive carcinoma (BRCA) (Figure 1D). These data support the development of a therapeutic intervention that can inhibit LAIR-1 checkpoint receptor interaction with collagen in the tumor microenvironment of solid cancers.

### NC410, a LAIR-2 human IgG1 fusion protein blocks collagen interaction with LAIR-1

Taking advantage of a natural decoy system that would both target cancers and reverse immune inhibition, we generated a LAIR-2 human IgG1 Fc fusion protein for therapeutic use named NC410 (Figure 2A). This fusion protein exists as a dimeric protein due to the cysteine bonding in the Fc portion of the protein. NC410 binds to collagen I and III with high avidity by Octet based binding studies and is cross-reactive to multiple species including rat and mouse due to the conserved nature of collagens across species (Figure 2B). Human LAIR-2 binds with much higher affinity to collagens than human LAIR-1, as reported elsewhere (42). NC410 completely blocks human LAIR-1 binding to collagen I, supporting the high avidity interaction of NC410 and its role as a LAIR-1 decoy therapeutic (Figure 2C). We also determined whether NC410 prevents LAIR-1 mediated signaling (Figure 2D, 2E). To do so, we used a reporter cell line that expresses the human LAIR-1 extracellular domain (ECD) fused to CD3z, thus conferring positive signaling capacity to LAIR-1 upon interaction with collagen ligands, and an NFAT-GFP reporter to visualize LAIR-1 mediated signal induction (21). Using flow cytometry (Figure 2D) and Incucyte microscope imaging (Figure 2E and Sup. Figure 1), we observed a dose-dependent inhibition of LAIR-1 NFAT-GFP reporter activity by NC410, indicating that NC410 inhibits collagen-mediated LAIR-1 signaling in a dose-dependent fashion.

**Figure 2.**
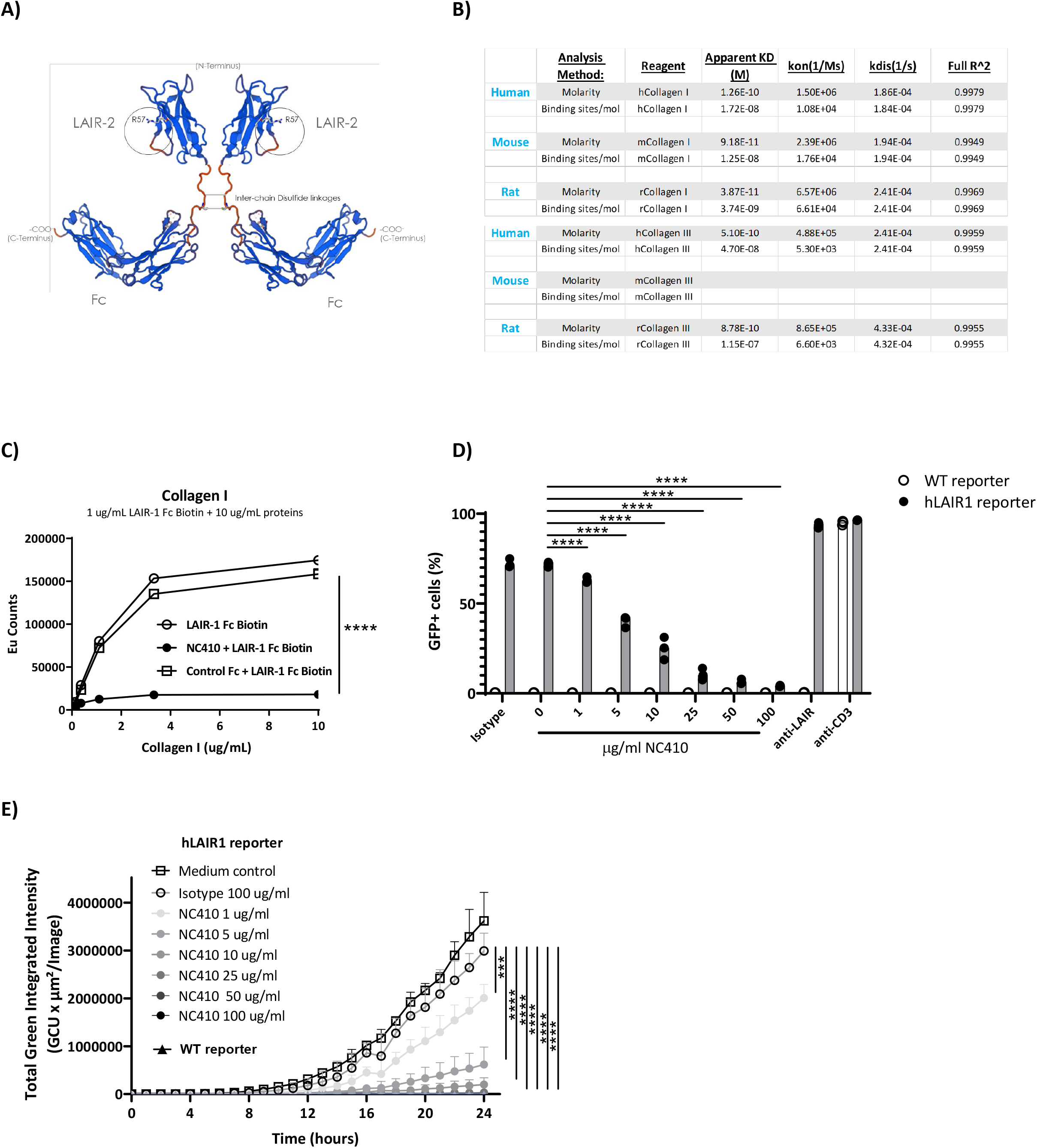
NC410 is an Fc dimeric fusion protein that blocks LAIR-1 interaction with collagen. A) NC410 is a biologic resulting by fusing LAIR-2 with a functional IgG1 to generate a dimeric fusion protein. B) Avidity characterization of NC410 to human, mouse and rat collagen I and III as measured by Octet analysis. C) Indicated amounts of collagen I were plate coated and the binding of soluble LAIR-1 was inhibited by NC410. Asterisks indicate statistical significance (****P < 0.0001, 2-way ANOVA). D and E) The LAIR-1 ECD was fused with CD3z and stably expressed in a cell line containing an NFAT-GFP pathway reporter. LAIR-1 ligation and CD3 ligation induce NFAT-GFP signaling. A parental cell line containing the CD3 NFAT-GFP reporter without LAIR-1 was used as control (WT). NC410 protein was added at increasing concentrations and inhibited collagen I (5ug/mL) mediated NFAT-GFP signaling through LAIR-1 binding by D) FACS analysis and E) Incucyte microscopy. Total green integrated intensity of WT and hLAIR-1 reporter cells is shown over time. Points represent the median of n=3 (with experimental triplicates in each independently performed experiment) and the whiskers indicate the 95% confidence interval (CI). Isotype control was used at the highest concentration (100ug/mL) and showed no inhibition of NFAT-GFP signaling. Anti-human LAIR (8A8 clone) and anti-mouse CD3 were used as positive controls. Closed circles in E) indicate NC410 treatment and open circles indicates control treatment. Significant differences between different treatment groups of hLAIR-1 reporter cells are indicated (and tested using a two-way ANOVA with Dunnett’s correction). In all plots: * P ≤ 0.05, ** P ≤ 0.01, *** P ≤ 0.001, **** P ≤ 0.0001.

### NC410 promotes T cell expansion and anti-tumor activity in humanized tumor models

To study the effect of NC410 on T cell function *in vivo*, we adoptively transferred human PBMCs into NSG mice. In this model, human xeno-reactive T cells against mouse antigens expand and cause xenogeneic-graft versus-host disease (xeno-GVHD). This model is often used to assess if targeting T cell stimulatory or inhibitory pathways alters xenogeneic T cell activity *in vivo*. We adapted this model by including the mouse tumor cell line P815 (Figure 3A). In this model, human PBMCs were injected intravenously on day 0, and P815 cells were injected subcutaneously a day later. NC410 or control Fc protein was administered on days 1 and 3 (Figure 3A). NC410 promoted the expansion of human CD8^+^ T cells on day 13, but not on day 6 in this model (Figure 3B). This was accompanied by reduced tumor growth of P815 after day 14 (Figure 3C).

**Figure 3.**
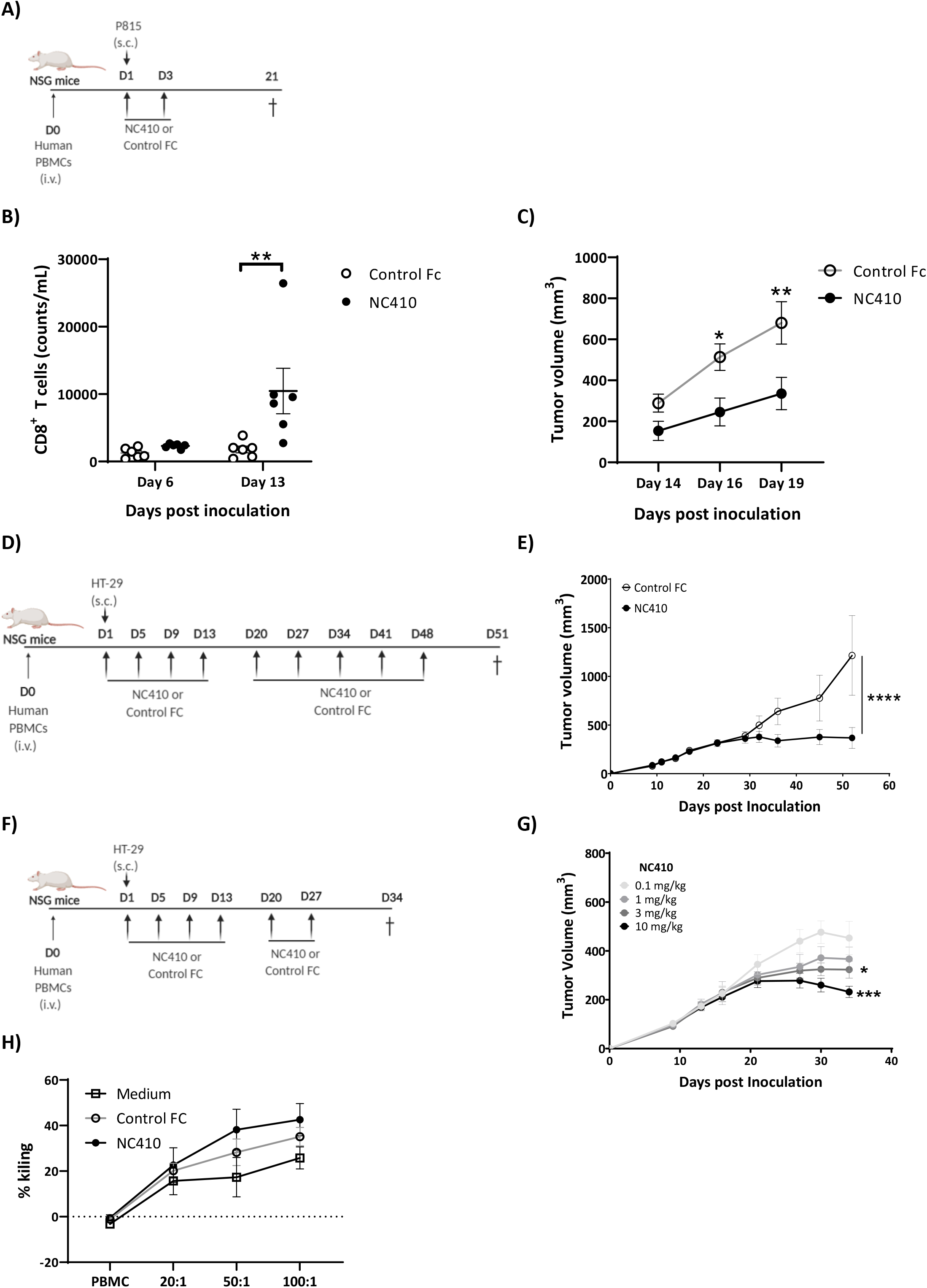
NC410 therapy promotes T cell expansion and tumor regression. A) Model for analysis of T cell expansion and xeno-anti-tumor response in the presence of NC410 or control Fc. 1.5 × 10^7^ total human PBMCs were adoptively transferred by intravenous injection to NSG mouse on Day 0. 2 × 10^5^ P815 tumor cells were injected subcutaneously on Day 1. Mice were treated on Days 1 and 3 with NC410 or control IgG1 (1 mg/kg) by intraperitoneal injection (N=6/group). B) CD8^+^ T cell expansion in blood on day 6 and 13. Asterisks indicate statistical significance (**P < 0.01, 2-way ANOVA with Sidak multiple comparisons). C) P815 tumor was measured every 2 - 3 days with a caliper and tumor volume was calculated. Asterisks indicate statistical significance (*P < 0.05, **P < 0.01, 2-way ANOVA with Sidak multiple comparisons). D) Humanized tumor model of HT-29 tumor injected subcutaneously in the presence of human PBMCs. 2 × 10^7^ total human PBMCs were adoptively transferred intraperitoneally (E) or intravenously (F) to NSG mice (N = 6/group) on Day 0. 1 × 10^6^ HT-29 tumor cells were injected subcutaneously with Matrigel on Day 1. Mice were treated with NC410 or control by intraperitoneal injection, Q4D x4 doses followed by Q7D until endpoint. Tumor growth was monitored 1 - 3 times a week E) Analysis of tumor growth in NC410 treated (10 mg/kg) vs control mice. Asterisks indicate statistical significance (**** P < 0.0001, 2-way ANOVA with Sidak multiple comparisons). F) Dose dependent effect of NC410. 6 mice per group were used for all experiments. Asterisks indicate statistical significance (*P < 0.05, ***P < 0.001, 2-way ANOVA with Tukey multiple comparisons). G) *In vitro* chromium release assay after 24h using HT-29 and PBMCs at 3 different effector to target ratios. 20:1 and 50:1 n=16 and 100:1 n=25 in 17 independently performed experiments. Closed circles indicate NC410 treatment and open circles indicates control treatment.

To determine if the T cell promoting and anti-tumor effects observed in the xenogeneic P815 model correlated with the capacity of NC410 to elicit T cell anti-tumor activity against an allogeneic human tumor, we developed a humanized subcutaneous tumor model using the HT-29 colorectal cancer cell line, which expresses collagens (Sup. Figure 2). In this model, NSG mice are injected intravenously with human PBMCs, and one day later subcutaneously with HT-29 cells (Figure 3C). NC410 or control treatments began on the same day as tumor implantation. NC410 treatment at 10mg/kg significantly reduced tumor growth compared to isotype control (Figure 3E). After titration of NC410, tumor volume was reduced in a dose dependent manner (Figure 3F). To investigate whether binding of NC410 to HT29 cells elicits antibody-dependent cellular cytotoxicity (ADCC), we performed *in vitro* cytotoxicity assays with HT-29 cells, using human PBMC as source of effector cells. We confirmed that HT-29 single cells expressed collagen and that NC410 was able to bind these cells after the EDTA treatment necessary to prepare the cells for the chromium release assay (Sup. Figure 3). In a short-term *in vitro* chromium release cytotoxicity assay, no differences in HT-29 killing by PBMC were observed between NC410, isotype treated samples and medium (Figure 3G) suggesting that enhancement of ADCC by NC410 is not contributing to tumor reduction *in vivo*. Thus, we established that NC410 has the capacity to induce T cell expansion and reduction of tumor growth in different *in vivo* tumor models.

### NC410 enhances anti-tumor T cell responses

Cytokines and chemokines mediate the host-response to cancer by activating and directing the trafficking of immune cells into the TME (43, 44). To determine if NC410 promoted infiltration and localized activity of T cells in the TME, immune profiling of HT-29 tumors and spleen tissues was performed at day 27 after treatment (10 mg/kg dose). HT-29 tumors were removed from euthanized mice, weighed, and dissociated for analysis of T cells and soluble factors. For equal comparison of systemic response, the spleen was harvested by identical means for analysis. Analysis of T cell numbers within the tumor showed a significant increase in the number of CD4^+^ and a trend of increased CD8^+^ T cells in tumors treated with NC410 compared to isotype control (Figure 4A and B). To determine the effector capacity of tumor infiltrating T cells (TILs) from NC410-treated mice, we re-stimulated TILs from digested tumors with anti-CD3+anti-CD28 for 5 hours and performed intracellular staining to examine IFN-γ and TNF-α production. After re-stimulation a significant increase in IFN-γ^+^ and IFN-γ^+^TNF-α^+^ double positive cells was observed in the NC410 treatment group compared to control. (Figure 4C). Based on this observation, we further assessed cytokines and chemokines in the local TME vs peripheral spleen on day 27 (Figure 4D-E and Sup figure 5). Analysis of cytokines indicated that IFN-γ and granzyme B were significantly increased in the TME, but not soluble CD40L (Figure 4D). Increased expression of CD40L and granzyme B was observed in the spleen (Figure 4D). Chemokine analysis indicated that CXCL10, 11 and 12 were all significantly increased in the TME, and significantly decreased in the spleen (Figure 4E). Importantly, the concentration of all three chemokines correlated with tumor reduction (Sup. Figure 4). These results support a role for NC410 in enhancing the recruitment of T cells and liberating their effector function in the TME.

**Figure 4.**
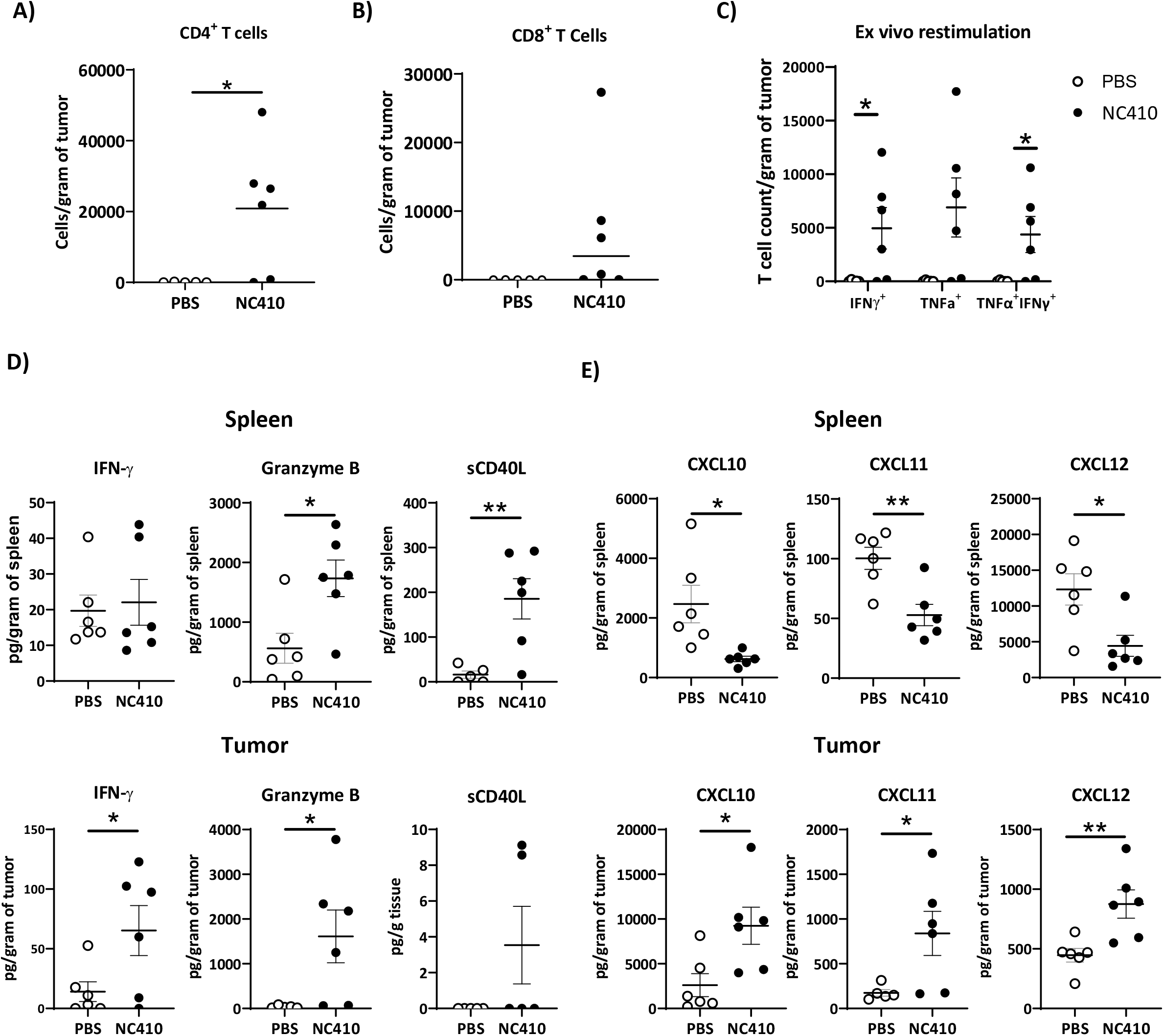
NC410 therapy induces immune activity in the local tumor microenvironment. 1 × 10^7^ total human PBMCs were adoptively transferred intravenously to NSG mice (N = 6/group) on Day 0. 1 × 10^6^ HT-29 tumor cells were injected subcutaneously with Matrigel on Day 1. Mice were treated with NC410 or control by intraperitoneal injection, Q4D x 4 doses followed by Q7D until endpoint. On day 27, tumor and spleen tissues were isolated for tumor infiltrating T cells and cytokine analysis. A) CD4^+^ and B) CD8^+^ TIL cell numbers in the tumor. The cell number was counted by flow cytometry and normalized to weight (gram) of tumor tissue. Asterisks indicate statistical significance (*P < 0.05, two-tailed t-test). C) Cytokine production by TILs following ex vivo restimulation with PMA and ionomycin for 5 hours. Cells were intracellularly stained for TNF-α and IFN-γ and the indicated cell populations were counted by flow cytometry. The data presents the cell counts normalized to the weight (gram) of the tumor tissue. Asterisks indicate statistical significance (*P < 0.05, two-tailed t-test). D-E) Analysis of tumor and spleen for cytokines (D) and chemokines (E) for analysis of local and systemic effects respectively. Tissue lysate protein was extracted from the tumor and spleen tissues. The cytokines were analyzed by Luminex and presented as the relative levels normalized to weight (gram) of tissue. Asterisks indicate statistical significance (*P < 0.05, **P < 0.01, two-tailed t-test).

### NC410 promotes collagen remodeling

Remodeling of ECM is pivotal to the development and progression of cancer. Recently it has been proposed that specific collagen-derived biomarkers reflecting the turnover of collagens may be used as tools to non-invasively interrogate cell reactivity in the TME and predict response to treatment (45–47). Because NC410 both engages collagens and promotes local T cell and immune responses, it was surmised that NC410 treatment may result in collagen remodeling. Therefore, we determined whether NC410 treatment increased collagen degradation products (CDPs) in the serum of HT-29 tumor bearing mice injected with PBMC over the course of tumor growth and rejection mediated by NC410 (Figure 5A and B). Nine CDPs were analyzed prior to experiment initiation and during four weeks of tumor growth (Figure 5C). While there was no change in most CDP, it was interesting that the concentration of two CDPs significantly increased in the serum of NC410 treated mice in comparison to controls. C6M, a collagen VI MMP-2 CDP, and C4GzB, a collagen IV Granzyme B CDP were significantly increased at week 4 (Figure 5D) in the NC410 treated group in comparison to the control group. Interestingly, this increase in serum CDPs was observed at the time of NC410 mediated tumor eradication suggesting that the CDPs were derived from the tumor and that the increase in CDPs was a result of T cells activation and effector function.

**Figure 5.**
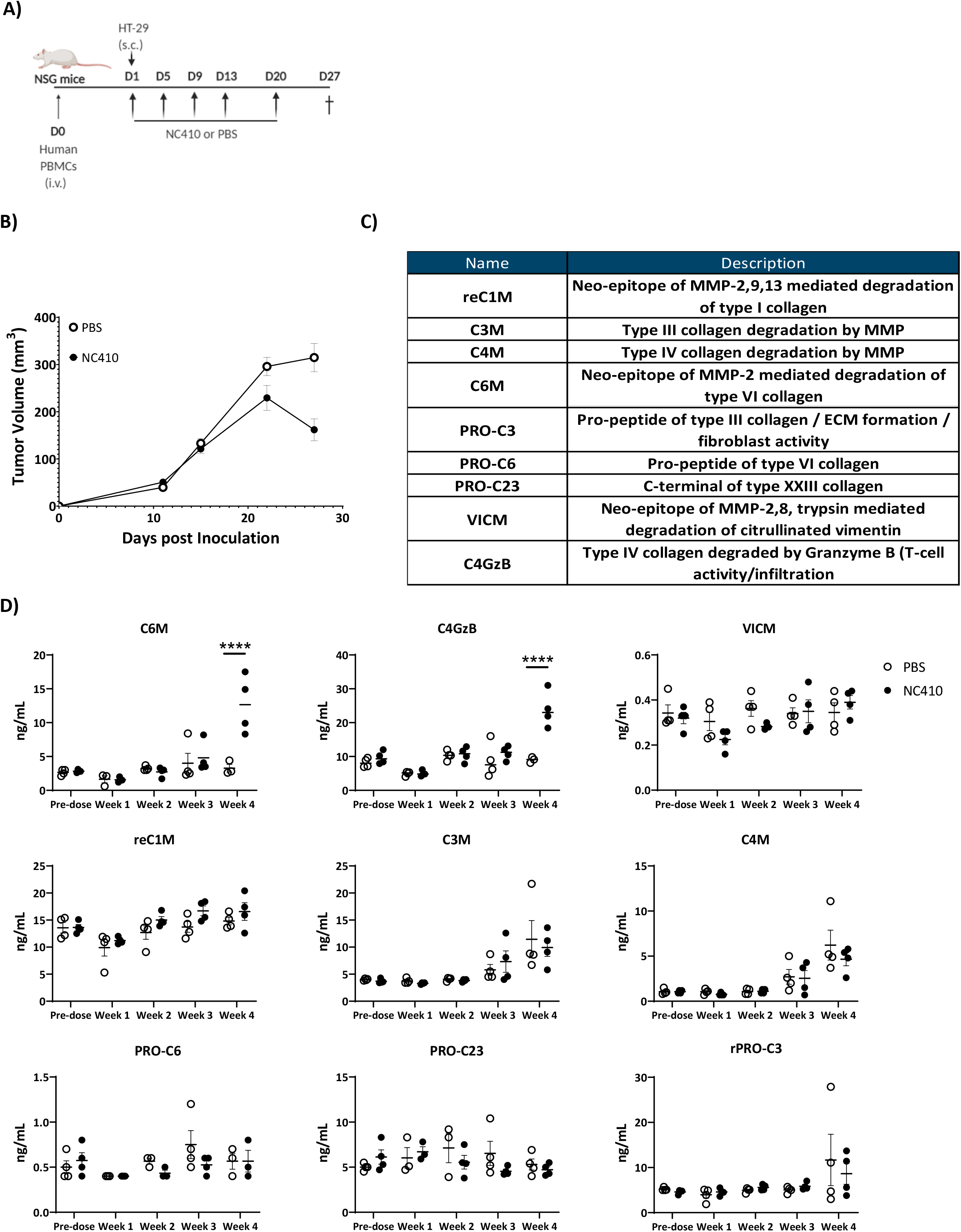
NC410 increases MMP2 and Granzyme B derived collagen fragments. A) Schematic representation of the humanized murine tumor model used. HT-29 tumor was injected subcutaneously in the presence of human PBMCs. Mice were treated with NC410 or control by intraperitoneal injection, Q4D x 4 doses followed by Q7D until endpoint. Mice were bled prior to start of experiment and weekly for 4 weeks. B) Tumor growth kinetics with NC410. Asterisks indicate statistical significance (*P < 0.05, **** P < 0.0001, 2-way ANOVA with Sidak multiple comparisons). C) List and description of collagen degradation products measured. D) Analysis of collagen degradation products in serum at baseline, at week 1, 2,3 and 4. MMP-2 derived collagen VI and granzyme B derived collagen IVI degradation products were increased at week 4 after NC410 administration. Closed circles indicate NC410 treatment and open circles indicates control treatment. Asterisks indicate statistical significance (**** P < 0.0001, 2-way ANOVA followed with Sidak multiple comparisons).

#### NC410 binds collagen rich tumors with an immune excluded phenotype

In order to identify tumor types and/or patient groups that would benefit from this therapy, we performed immunohistochemical analysis of serial tissue sections from seven different tumor types, of which we analyzed biopsies from ten patients per tumor type. Sections of head and neck squamous cell carcinoma (HNSC), glioblastoma (GBM), melanoma, non-small-cell lung carcinoma (NSCLC), high-grade serous carcinoma (HGSC), pancreatic ductal adenocarcinoma (PDAC), and stomach adenocarcinoma (STAD), were stained for hematoxylin and eosin, Masson’s Trichome, anti-LAIR-1, NC410 and the immune cell markers, CD45, CD3, CD68 and CD163 (Figure 6A). NC410 binding co-localized with Masson’s Trichrome positive collagen areas within the TME and LAIR-1^+^ immune cells were present in all tumors (Figure 6B, Sup. Figure 5, heathy tissue Sup. Figure 6). Both myeloid and lymphoid cells within the TME expressed LAIR-1 (Sup. Figure 7). NC410 binding was highest in pancreatic cancer in agreement with a collagen-rich microenvironment (Figure 6C). Head and neck squamous cell carcinoma had the highest number of LAIR-1^+^ cells (Figure 6C). Importantly, LAIR-1^+^ cells were enriched in NC410 positive areas (Figure 6D) suggesting that LAIR-1^+^ cells were entrapped in the collagen matrix. Most human solid tumours exhibit distinct immunological phenotypes being divided into immune inflamed, immune excluded, and immune desert tumors on the basis of immune cell infiltrate and localization (48, 49). By characterizing our cohort according to these immunological phenotypes (Figure 6E, Sup. Figure 8), we observed that NC410 binding was particularly increased in immune excluded tumors, namely HNSC, melanoma, NSCLC and STAD. (Figure 6F). Thus, in immune excluded tumors, LAIR-1^+^ cells are sequestered in collagen-rich areas where NC410 can bind.

**Figure 6.**
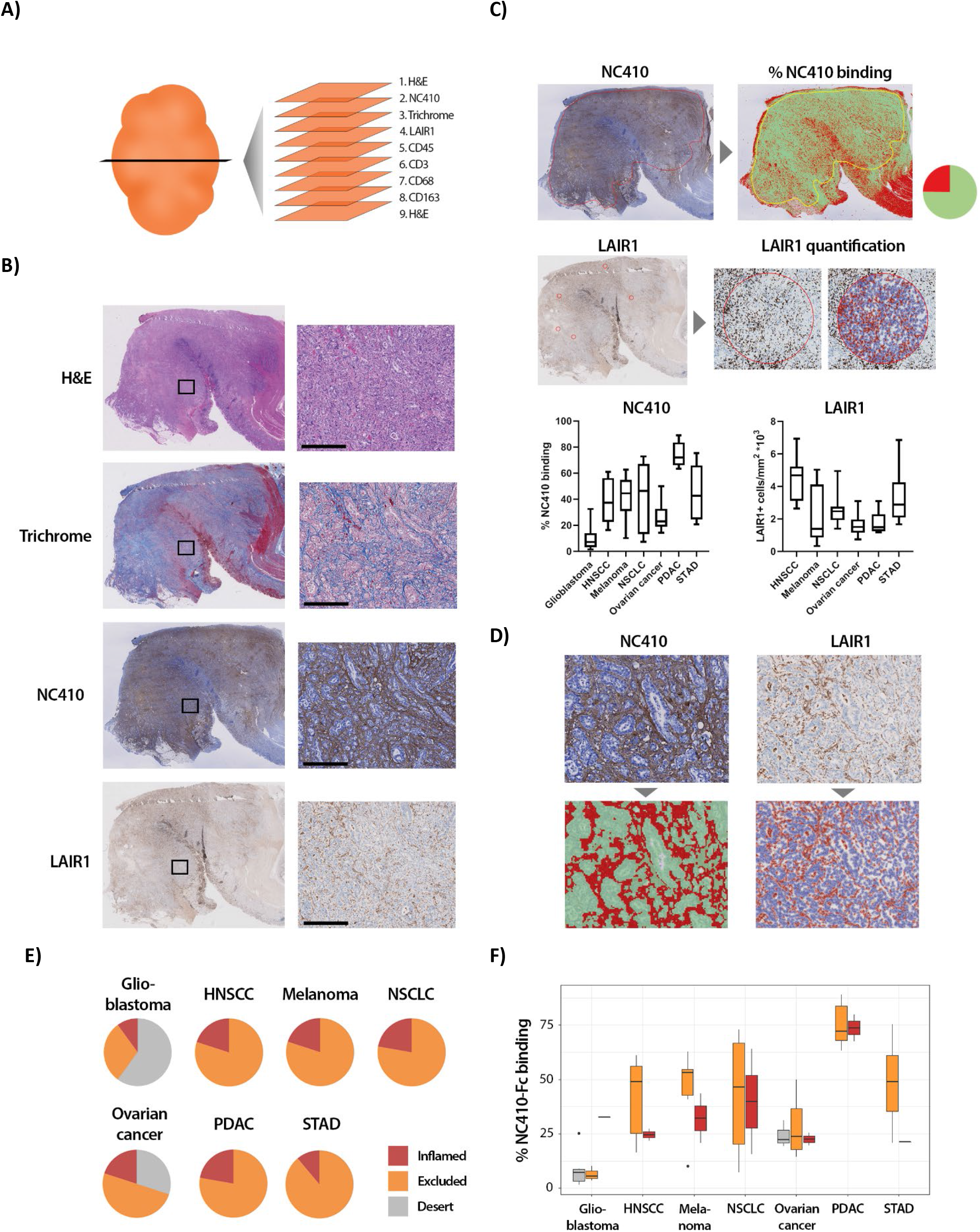
NC410 binds to collagen rich tumors with an immune excluded phenotype. A) Schematic representation of the immunohistochemical stainings performed. B) Representative hematoxylin and eosin (H&E), Masson Trichrome, NC410 and LAIR-1 staining in a stomach tumor specimen. C) Quantification of LAIR-1 and NC410 staining of 10 patients per tumor type across 7 different tumor types (Head and neck (HNSC), melanoma, non-small cell lung carcinoma (NSCLC), ovarian, pancreatic (PDAC) and stomach cancer (STAD)). In the associated piecharts, we show the percentage of NC410 binding where green indicates the percentage of NC410 positive area and red indicates the percentage of NC410 negative area. D) Higher magnification pictures of stomach cancer specimens show LAIR-1^+^ cells (depicted in red, right side) colocalizing with NC410 positive areas (depicted in red, left side). E) Patients within each tumor type were characterized into immune desert, immune excluded and immune inflamed based on CD3^+^ T cell presence and localization. F) Quantification of NC410 staining across the different immune phenotypes in the 7 different tumor types analyzed.

## Discussion

Cancer cells have evolved multiple mechanisms to escape immune surveillance, such as defects in antigen presentation machinery, the recruitment of immunosuppressive cell populations and the upregulation of negative regulatory pathways, all resulting in hampered effector function of immune cells and the abrogation of antitumor immune responses (50). Tumor progression is also accompanied by extensive remodeling of the extracellular matrix leading to the formation of a tumor-specific ECM, which is often more collagen-rich and of increased stiffness (51, 52). Collagen expression and density have been shown to be associated with a worse prognosis, either by directly promoting tumor growth or by preventing immune cell infiltration (8).

We have shown that a combined collagen signature was associated with a worse prognosis in 13 out of 21 tumor types supporting the notion that increased collagen expression is detrimental for overall survival in tumors. Importantly, LAIR-1 mRNA expression was associated with a worse prognosis in 4 out of 21 tumors and LAIR-2 mRNA expression was associated with a better prognosis in 6 out of 21 tumors. This indicates that targeting LAIR-1-collagen interaction might be helpful in a specific group of tumors.

It has been demonstrated that collagens can also directly modulate T cell function(53–55), for instance through the inhibitory collagen receptor LAIR-1(18, 27). We hypothesize that increased expression and remodeling of collagen in the TME serves to set a threshold of T cell activation through LAIR-1 and is employed by tumor cells to escape immune surveillance. In this study we show that disrupting collagen interaction with LAIR-1 by means of a dimeric LAIR-2 Fc fusion protein has a therapeutic effect in cancer models. NC410 administration in humanized mouse tumor models results in a reduction of tumor volume by increasing T cell expansion and cytotoxic activity but does not enhance ADCC *in vitro*. NC410 binds to collagen-rich areas of tumors, where LAIR-1^+^ cells are sequestered. NC410 also binds to healthy collagen, therefore next to the direct effect on T cell mobilization and activation in the tumor, systemic effects could also add to its therapeutic potential.

During ECM turnover, proteolytically cleaved matrix degradation fragments are released into the systemic circulation (3). These small protein fragments containing specific protease-generated neo-epitopes, or ‘protein fingerprints’, can be used as serological biomarkers directly reflecting disease pathogenesis. Several studies have shown that serum levels of collagen degradation fragments are elevated in cancer patients compared to healthy controls (56–58), and that checkpoint blockade may alter serum CDP levels (47). We observed that specific fragments generated by degradation of collagen VI by MMP-2 (C6M) and collagen IV by granzyme B (C4GzB) were increased after NC410 treatment. The increase of circulating C6M and C4GzB coincided with reduction of tumor volume suggesting that these fragments might be generated by increased infiltration and/or activation of T cells in the TME. C6M and C4GzB collagen fragments have the potential to be biomarkers of NC410 response, which will be investigated in a recently initiated NC410 first-in-human clinical trial (NCT04408599).

In the era of personalized medicine, it is very important to address how to select patients that are likely to benefit from immune checkpoint blockade therapy. Within different tumor types a distinction based on the presence and location of immune infiltrates can be made, separating patients into having immune excluded, immune inflamed and immune desert tumors. Currently, immune inflamed tumors, where the majority of immune cells are present throughout the neoplastic cells, have been associated with better response to currently available checkpoint blockade therapy compared with the other phenotypes (59, 60). The immune-excluded phenotype is characterized by the presence of immune cells that cannot penetrate the parenchyma of the tumors but instead are located in the stroma that surrounds the cancer cells (61). These tumors are generally resistant to current checkpoint blockade therapy (62). We performed an immunohistochemical screen across 7 tumor types, analyzing biopsies of 10 individual patients per tumor type, and observed that immune excluded tumors had the highest NC410 binding. Importantly we observed that LAIR-1^+^ cells were present within these NC410-binding collagen rich areas. This might indicate that an abundance of collagen keeps the immune cells trapped in the tumor stroma, possibly by binding to LAIR-1. Treatment with NC410 could block this interaction resulting in LAIR-1^+^ cell activation and infiltration into the tumor nests, promoting tumor clearance.

LAIR-2 acts as a decoy receptor by binding to collagen with higher affinity than LAIR-1 and therefore antagonizes LAIR-1 inhibitory function (33, 34). Circulating LAIR-2 protein concentration is low or undetectable in healthy individuals (33, 42). In the presence of limited endogenous LAIR-2, LAIR-1 can bind collagens, thereby allowing the inhibitory receptor to signal and prevent or reduce T cell activity and/or retain LAIR-1+ cells in collagen-rich areas. In the TME increased levels of collagens, in absence of increased LAIR-2, will therefore promote tumor immune escape. Our analysis of TCGA data indeed revealed that enhanced expression of endogenous LAIR-2 in some cancers associated with better prognosis, suggesting that further increasing LAIR-2 in vivo would have a therapeutic advantage. NC410 has a higher avidity to collagen than endogenous LAIR-2 due to its dimeric structure, since *in vivo* LAIR-2 is expressed as a monomer, enhancing its potential to block the inhibitory capacity of membrane-bound LAIR-1. NC410 binds both healthy and tumoral collagen and theoretically could be sequestered before reaching the tumor site. We hypothesize that the avidity of NC410 towards tumoral collagen is higher than to healthy collagen, therefore resulting in a therapeutic effect, but this needs additional study. Other LAIR-1 ligands, such as C1q have been shown to be increased in tumors (19) and may also provide a local inhibitory effect that could be potentially reversed by NC410.

Taken together, tumor patients with collagen rich tumors that present an immune excluded phenotype and low endogenous LAIR-2 expression would be predicted to benefit the most from NC410 therapy. Our immunohistochemical analysis points to HNSC, melanoma, NSCLC and STAD as immune excluded tumor types with the highest NC410 binding. In agreement with our hypothesis that blocking LAIR-1-collagen interaction in these tumors would be beneficial, TCGA analysis showed that increased LAIR-2 mRNA at the tumor site also increases survival probability in HNSC, melanoma, and STAD. Together, this may be indicative that patients suffering from these particular tumors might benefit the most from NC410 treatment.

While conducting our study, an unrelated research group generated a structurally similar LAIR-2 Fc fusion protein for use as an immune therapy for cancer to block LAIR-1 inhibitory function (63). Xu *et al.* showed that LAIR-2 Fc reversed T cell inhibition *in vitro* and *in vivo* and promoted anti-tumor immunity *in vivo*. Our models are non-overlapping with the models described in that study, independently supporting the concept of LAIR-2 Fc for cancer immunotherapy. In addition, we now provide an immunohistochemical rationale for LAIR-2 Fc treatment in certain tumor types by extensive analysis of human tumor samples, as well as potential biomarkers for patient selection and response to therapy.

Our data support NC410 as a novel immunomedicine for targeting immune excluded collagen-rich tumors and enabling normalization of the tumor-immune microenvironment. First-in-human studies have recently been initiated with NC410 (NCT04408599).

## Material and Methods

### Mice

NSG (NOD-SCID IL2Rγnull) female mice were purchased at the age of 6 - 8 weeks old from Jackson Labs. Upon arrival at NextCure, mice were divided into 5 - 6 mice per cage and kept in the quarantine room for at least 6 days to acclimate to the environment. All mouse studies were performed at NextCure based on Institutional Animal Care and Use Committee standards.

### Bioinformatics

For collagen expression, the mRNA expression of 44 collagen genes in normal and tumor tissue for each cancer type were queried using TCGA database (https://cancergenome.nih.gov/). mRNA collagen data was available in TCGA except for COL6A5. The transcript per million (TPM) values were log2 transformed and averaged per individual across all collagen proteins. The distribution of average expression across individuals was plotted using ggplot utility version 2.3.3.2 in R version 4.0.2. The distribution of expression in normal tissues was compared against those in cancer tissue using a non-parametric Wilcoxon test in R.

For overall survival analysis, the log2 transformed collagen expression for each of the 43 collagen genes in various cancers was divided into four quantiles. The patients in lower quantile were considered individuals with low expression and those in the upper quantile were considered those with high expression. The estimate of survival based on collagen expression in the tumor was determined using the Kaplan-Meier method. The survival curves were drawn using ggsurvplot function in the survminer R package. The survival curves for the protein-cancer combination where the p-value was less than 0.05 were considered significant. The expression and overall survival analyses for LAIR-1 and LAIR-2 were performed in the same manner. R codes can be found in the source code files.

For collagen gene expression analysis of HT-29 cells, data was acquired from the public dataset GSE41586 (https://pubmed.ncbi.nlm.nih.gov/23902433/). Raw count data of untreated HT-29 cells was retrieved and normalized using the DESeq2 package (v1.28.1) in R (v4.0.2). Data was then log2 transformed and plotted using the ggplot package (v3.3.2).

### Cells and antibodies

2B4 T cell hybridoma cells transduced with a NFAT-GFP reporter and hLAIR-1-CD3ζ, the hLAIR-1 reporter cells, or transduced with a NFAT-GFP reporter and CD3ζ, the WT reporter cells (21), were cultured in RPMI 1640 (Life Technologies) supplemented with 10% Fetal Bovine Serum (FBS) (Sigma-Aldrich) and 1% penicillin/streptomycin (Gibco). P815 cells (ATCC) were cultured in DMEM, 2 mM L-glutamine, 25 mM HEPES, 10% FBS and Penicillin-Streptomycin (100 U/mL-100 μg/mL). HT-29 cells (ATCC) were cultured in IMDM, 2 mM L-glutamine, 25 mM HEPES, 10% FBS and Penicillin-Streptomycin (100 U/mL-100 μg/mL). CHO cells were cultured in CD CHO medium (ThermoFisher). Anti-human CD45 (hCD45)-BV421, anti-mouse CD45(mCD45)-APC, anti-human CD3(hCD3)-Percp.Cy5.5, anti-human CD8(hCD8)-AF488, anti-human TNF-α (hTNF-α)-PE and anti-human IFN-ɣ (hIFN-ɣ)-PECy7 were from ThermoFisher. Anti-human CD4 (hCD4)-BV711 was from Biolegend.

### LAIR-1-Fc and LAIR-2-Fc (NC410) generation

The human LAIR-1 and LAIR-2 genes were synthesized by GeneArt and genetically fused with the N terminus of IgG1 Fc domain. A stable CHO cell line expressing recombinant human LAIR-1 Fc or LAIR-2 Fc fusion protein was developed using the Lonza GS system. Briefly, 5 × 10^7^ CHO cells were transfected by electroporation using 80 μg of linearized plasmid DNA in a 0.4 cm cuvette. Following electroporation (300 V, 900 μF) cells were resuspended in 100 mL glutamine-free CD CHO medium (ThermoFisher). The following day, MSX (Millipore) was added to a final concentration of 50 μM, and cells were monitored for the next two weeks as prototrophic cells began to grow. Single clone of stably transfected cells was cultured, and supernatant was harvested and purified by affinity chromatography. The protein purity was determined by HPLC and sodium dodecyl sulfate-polyacrylamide gel electrophoresis. For the LAIR-1-collagen binding assay, LAIR-1 Fc was biotinylated with EZ-Link NHS-PEG4-Biotin (ThermoFisher), No-Weight Format (ThermoFisher) and free biotin was removed by ZebaSpin Desalting Columns (ThermoFisher) following manufacturer’s instructions.

### Human PBMC preparation for *in vitro* experiments

Peripheral Blood Mononuclear Cells (PBMCs) were isolated from blood of healthy donors (in agreement with ethical committee of the University Medical Center Utrecht (UMCU) and after written informed consent from the subjects in accordance with the Declaration of Helsinki) using standard Ficoll density gradient centrifugation. Briefly, blood was diluted 1:1 with PBS and layered on top of 15 mL of Ficoll-Paque (GE healthcare) in 50 mL conical tube. Suspension was centrifuged at 400 × g for 20 min at 20 °C in a swinging bucket rotor without brake. The mononuclear cell layer at the interphase was carefully collected and transferred to a new 50 mL conical tube. The cells were washed with PBS and centrifuged at 300 × g for 10 min at 20 °C. The supernatant was discarded, and the cell pellet was washed twice with 50 mL PBS. The isolated PBMCs were immediately used for *in vitro* studies.

### Binding and Blocking Studies

#### Octet avidity analysis

A ForteBio Octet RED96 instrument was used for avidity assessments. The anti-human Fc antibody capture sensors (ForteBio) were first loaded with LAIR-2-Fc followed by an association step where the loaded sensor was dipped into wells containing human, mouse or rat collagen I (human, R&D Systems; mouse, Ray Biotech; rat, Yo Protein) or collagen III (human, R&D Systems; mouse, Abbexa; rat, Yo Protein). NC410 protein was diluted in assay buffer (ForteBio) at 20 μg/mL and the collagen concentration ranged from 1.56 μg/mL to 100 μg/mL for collagen I and from 0.78 μg/mL to 50 μg/mL for collagen III. Data processing was conducted using the Octet’s Data Analysis 9.0 software.

#### Time Resolved Fluorometry (TRF) Immunoassay

EIA plates were coated with human collagen I (StemCell) in 0.01 N HCL (100 μL/well) overnight at 4 °C. The following day, plates were equilibrated to ambient temperature and washed 3 times (300 μL/well) with DELFIA Wash buffer (PerkinElmer). The plates were blocked for non-specific binding with 3% BSA (200 μL/well, Millipore) for 1 hour. Plates were washed 3 times (300 μL/well) with DELFIA wash buffer and an NC410-biotin and human LAIR-1 Fc mixture (50 μL/well) was added to plates and incubated for 2 hours at ambient temperature. The plates were washed 3 times (300 μL/well) with DELFIA wash buffer. Europium-labeled Streptavidin (Eu-SA) (100 μL/well, PerkinElmer) was diluted 1:1000 in DELFIA assay buffer and was added to plates and incubated for 1 hour at ambient temperature. Following the incubation, the plates were washed with 300 μL/well of DELFIA wash buffer. DELFIA enhancement solution was equilibrated to ambient solution during detection antibody incubation. Following the last wash, 100 μL of DELFIA enhancement solution (PerkinElmer) was added to each well and incubated on a plate shaker for 5 minutes prior to reading on an EnVision plate reader with excitation at 340 nm, and fluorescence reading at 615 nm (PerkinElmer).

#### Flow cytometry blocking studies

To assess the blocking capacity of NC410 a titration assay with LAIR-1 reporter cells was performed as previously described(21). Black Falcon clear flat bottom 96-well plates were coated with 5 μg/mL human collagen I (Sigma-Aldrich) in 2 mM acetic acid (Merck), anti-mouse-CD3 (BD), anti-human-LAIR-1 antibody (clone 8A8 (21)) in PBS (Sigma-Aldrich) or isotype control (eBiosciences) in PBS by spinning down for three minutes at 1700 rpm and incubating overnight at 4 °C. The next day, plates were washed with PBS and pre-incubated with the indicated concentrations of NC410 or isotype control (Nextcure) in culture medium by spinning down for five minutes at 1500 rpm at room temperature (RT) and incubating for two hours at 37 °C.

WT and hLAIR-1 reporter cells were harvested and seeded at 1 × 10^6^ cells/mL in 50 μl/well on top of the collagen and fusion proteins treated wells and spun down for three minutes at 1700 rpm at RT. Plates were incubated overnight, approximately 16 hours, at 37 °C and GFP expression was measured on a LSRFortessa (BD Biosciences).

#### Incucyte assay

Plates were prepared similarly to the flow cytometric analysis. After adding reporter cells to collagen and fusion protein treated wells, plates were placed in the Incucyte S3 (Sartorius) and green fluorescence of the GFP expressed by the reporter cells was imaged every hour for 24 hours.

Analysis of Incucyte images was performed using the Incucyte 2020A analysis programme (Sartorius), where green fluorescence was evaluated using Top-Hat segmentation (radius 100 μm and threshold 2 GCU), edge split turned on, minimum mean intensity of 3 GCU and an area filter of 600 μm^2^ to calculate the total green integrated intensity (GCU x μm^2^/Image) per well.

### *In vivo* experiments

Leukopaks (StemCell) were diluted with PBS and layered with 35 mL of diluted cell suspension over 15 mL of Ficoll-Paque (GE healthcare) in 50 mL conical tube. Suspension was centrifuged at 400 × g for 30 min at 20 °C in a swinging bucket rotor without brake. The mononuclear cell layer at the interphase was carefully collected and transferred to a new 50 mL conical tube. The cells were washed with PBS and centrifuged at 300 × g for 10 min at 20 °C. The supernatant was discarded, and the cell pellet was washed twice with 50 mL PBS. The isolated PBMCs were frozen and stored in liquid nitrogen.

Prior to *in vivo* studies, PBMCs were rested overnight in RPMI 1640, 2 mM L-glutamine, 10 mM HEPES, 10% FBS, Penicillin-Streptomycin (100 U/mL-100 μg/mL) and 250 U/mL DNase (Millipore). Female NSG mice were injected intraveneously (i.v.) with 1-2 × 10^7^ PBMC in 100 μl of 1x PBS. The next day, 2 × 10^5^ P815 cells in PBS, or 1 × 10^6^ HT-29 cells in PBS with 50% Matrigel (Corning) were injected subcutaneously on the right flank. Mice were randomly assigned into treatment or control groups (6 mice per group). The sample size per group was determined with resource equation approach n=DF/k +1 where n=number of sample per group, k=number of groups and DF=degrees of freedom with acceptable range between 10 to 20 in analysis of ANOVA and t test(64, 65). Beginning on day 1, LAIR-2 Fc and control protein were injected intraperitoneally (i.p.) Q4D x 4 doses followed by Q7D until the endpoint. Tumor size was monitored 2-3 times a week. Tumor volumes were determined according to the formula tumor volume = 0.5 × (shorter diameter)^2^ × longer diameter. At endpoint, tumor and spleen tissues were collected for T cell population and/or cytokine analysis. In some studies, blood was collected weekly for T cell population and collagen degradation analysis. Mice were randomized in all tumor models prior to treatment. Tumor measurements were performed in a blinded manner.

For ex vivo analysis, cells were stained with Zombie NIR viability dye (Biolegend) in PBS at RT for 10 min. After washing with FACS buffer (2%FBS in PBS), cells were stained with antibodies against cell surface antigens at 4 °C for 30 min. For the intracellular staining, cells were stimulated with Cell Stimulation Cocktail plus protein transport inhibitors (ThermoFisher) at 37 °C for 5 hours followed by Zombie NIR and cell surface antigen staining. After cell fixation and permeabilization, the intracellular TNF-α and IFN-γ were stained following the instructions of BD Cytofix/CytoPerm Plus Fixation/Permeabilization Kit. All antibodies were used at the concentrations recommended by manufacturers. Stained cells were washed and resuspended in 150 μL FACS buffer and 80μL of samples were acquired on an Attune flow cytometer (ThermoFisher). Human CD4^+^ and CD8^+^ T cells were gated based on live/hCD45^+^mCD45^−^hCD3^+^hCD4^+^hCD8^−^ and live/hCD45^+^mCD45^−^hCD3^+^hCD4^−^hCD8^+^ respectively.

To prepare mouse blood T cells for staining, 80 – 200 μL blood was treated with 3 mL ACK lysis buffer (KD medical) to lyse the red blood cells for 5 min at RT followed by washing with PBS. The initial volume of blood was recorded for calculation of cell counts per mL of blood according to the formula: Cell counts per mL of blood= [acquired counts × 150 × 1000] ÷ [initial blood volume(μL) × 80].

To prepare the single cell population from tumors for staining, tumor tissues were weighed, cut into small pieces, and digested with mouse tumor dissociation kit (Miltenyi) and dissociated with gentleMACS Dissociator (Miltenyi). The tumor weight was recorded for the normalization of cell counts.

To assess cytokines from tissues, tumor and spleen tissues were weighed and cut into small pieces in 1.5mL Eppendorf tube on ice. 200 μL of RIPA Lysis buffer (ThermoFisher) was added with proteinase inhibitor (Roche) and 250 U/mL of DNase (Millipore) and the tissues were dissociated with pellet pestles (Sigma) on ice. The samples were kept on ice for 30 min, vortexing occasionally. Centrifuge at 10,000 ×g for 20 mins at 4 °C to pellet cell debris and then transfer the supernatant to a fresh Eppendorf tube without disturbing the pellet. The tissue weight was recorded for the normalization analysis of cytokines.

### Luminex cytokine assay

Tissue lysate was used for analysis of the cytokine profile (SDF-1α, IL-2, IL-4, IP-10, IL-10, IL-17A, IFN-γ, TNF-α, I-TAC, granzyme B, sCD40L) using a Luminex assay. The Singleplex Luminex™ Protein Assay Kit for each cytokine was from ThermoFisher. Antibody-specific capture magnetic beads were added to wells of a 96-well plate. Samples and protein standards were then placed into the microplate wells. After incubation, the beads were washed using a handheld magnet and were resuspended in secondary detection antibody solution followed by washing and addition of streptavidin-RPE. The beads were then washed again for analysis on a FlexMAP 3D™ (ThermoFisher).

### EDTA treatment of HT-29 cells

HT-29 cells were collected from T75 flasks (Thermo Scientific) using different concentrations of EDTA (0.1; 0.2; 0.5; 1 and 2 mM) for 10 min at 37C. Cells were blocked with 10% BSA/10% normal mouse serum (NMS)/ 10%FCS for 15 min at 4C. Cells were then incubated with biotin labelled NC410 (10ug/ml; Nextcure) in PBS + 1% BSA buffer for 30 min RT. After washing with PBS + 1% BSA, by spinning down for five minutes at 1500 rpm at RT, cells were incubated with streptavidin APC (eBioscience) diluted in PBS + 1% BSA buffer for 20 min at 4C. Cells were washed again and measured on a FACSCanto (BD biosciences).

### Collagenase treatment of HT-29 cells

HT-29 cells were collected from T75 flasks (Thermo Scientific) using 0.1mM EDTA for 10 min at 37C. Cells were washed and treated with 40 U Collagenase (from Clostridium histolyticum; Sigma) for different time points (5, 10, 15, 20 and 30 min) at 37C. Cells were washed, by spinning down for five minutes at 1500 rpm at RT, and blocked with 10% BSA/10% normal mouse serum (NMS)/ 10%FCS for 15 min at 4C. Cells were washed and incubated with biotin labelled NC410 (10ug/ml; Nextcure) in PBS + 1% BSA buffer for 30 min RT. After washing with PBS + 1% BSA, by spinning down for five minutes at 1500 rpm at RT, cells were incubated with streptavidin APC (eBioscience) diluted in PBS + 1% BSA buffer for 20 min at 4C. Cells were washed again and measured on a FACSCanto (BD biosciences).

### Antibody dependent cellular cytotoxicity (ADCC) Assay

ADCC with ^51^Cr-labeled target cells was described previously (66). Briefly, HT-29 target cells were labeled with 100 μCi (3.7 MBq) ^51^Cr for 3 h in complete medium. After extensive washing, cells were adjusted to 10^5^/mL. HT-29 cells were then incubated with NC410 or isotype control for 30 min. Different effector to target ratios (E:T) were made by adding increasing amounts of PBMCs to NC410 or isotype treated HT-29 cells per well (a fixed amount of 10.000 tumor cells was used) in round-bottom microtiter plates (Corning). After 24 h of incubation at 37°C, ^51^Cr release was measured in counts per minute (cpm). The percentage of specific lysis was calculated using the following formula: % lysis = [(counts of sample–minimum release)/(maximum release–minimum release)] × 100. Target cells with PBMCs in complete medium or supplemented with 5% Triton X-100 (Roche Diagnostics) were used to determine minimum and maximum release, respectively.

### Immunofluorescence staining

5000 HT-29 cells were seeded in black Falcon clear flat bottom 96-well plates and cultured for 3 days. Cells were fixed with 4% paraformaldehyde for 15 min at room temperature, and blocked with 5% BSA in PBS for 1 h at room temperature. Cells were then incubated with isotype control (Nextcure), biotin labelled NC410 (10ug/ml, Nextcure) or pan-collagen antibody (Thermo fisher) diluted in PBS + 1% BSA buffer for 1 h at room temperature. After thoroughly washing with PBS, the slides were incubated with anti-human IgG1-AlexaFluor 594 or streptavidin Alexa Fluor 594 (Life Technologies-Thermo Fisher Scientific) diluted in PBS + 1% BSA buffer for 30 min at room temperature. Slides were finally washed and mounted with DAPI VectaShield hardset (Vector Lab) and allowed to settle before image acquisition on a Zeiss fluorescence microscopy (Zeiss) using the Axiovision software (Zeiss).

### Collagen degradation peptide analysis

The mouse serum samples were assessed for collagen degraded peptides (PRO-C3, C3M, C4M, G4GzB, RRO-C6, VICM, CRPM, reCIM) by Nordic Bioscience as previously described (47, 67).

### Tumor specimens for immunohistochemistry

Specimens of seven selected tumor types were included for analysis: head and neck squamous cell carcinoma (HNSC), glioblastoma (GBM), melanoma, non-small-cell lung carcinoma (NSCLC), high-grade serous carcinoma (HGSC), pancreatic ductal adenocarcinoma (PDAC), and stomach adenocarcinoma (STAD). Of each tumor type, in agreement with the ethical committee of the UMCU, formalin fixed, paraffin embedded (FFPE) material of 9-10 tumor specimens and five healthy specimens was collected from the tissue biobank (research protocol 17-786). The cohort of HNSC tumors comprised tongue tumors with a diameter of 10 mm or larger. Only melanomas with a Breslow thickness of 0.5 mm or higher were included. Tissue was obtained from primary tumors. Patients did not receive any systemic treatment or radiotherapy before the tumor specimens were obtained. Healthy specimens were preferably separate tissue blocks obtained from the same patients the tumor specimens were obtained from. In case of glioblastoma, healthy specimens were collected post-mortem.

### Immunohistochemistry

The following stainings were performed on consecutive slides of all tumor specimens: H&E, Masson’s Trichrome, LAIR-1, NC410, CD45, CD3, CD68, and CD163. The LAIR-1, CD45, CD3, CD68, and CD163 stainings were performed using a Ventana Bench Mark XT Autostainer (Ventana Medical Systems, Tucson, AZ, USA). Stainings were performed on 4 μm sections of each tissue block. NC410 was biotinylated and used for manual, immunohistochemical staining. Antigen retrieval was performed by incubating the slides at 100°C for 24 or 64 mins in either EDTA or Citrate buffer, as indicated in the table below. For all assays, the sections were incubated with the primary antibody for 1 hour at RT. Subsequently, the sections were incubated with HRP-labeled secondary antibody, developed using H2O2 and DAB and counterstained with Hematoxylin.

| Antibody | Supplier | Clone | Lot no | Pretreatment | Dilution |
| --- | --- | --- | --- | --- | --- |
| LAIR-1<br>(polyclonal) | ATLAS<br>antibodies | - | A103834 | 24' Citrate pH6.0 | 1:500 |
| LAIR-1<br>(monoclonal) | BD<br>Pharmingen | DX26 | 550810 | 64' EDTA pH9.0 | 1:100 |
| NC410 | Nextcure | - | - | 24' Citrate pH6.0 | 1:500 |
| CD45 | DAKO | PD7/26+2B11 | 110 | 24' EDTA pH9.0 | 1:25 |
| CD3 | DAKO | - | 20042625 | 24' EDTA pH9.0 | 1:100 |
| CD68 | Novocastra | KP1 | 6009203 | 24' EDTA pH9.0 | 1:1600 |
| CD163 | Novocastra | 10D6 | 6064616 | 24' EDTA pH9.0 | 1:800 |

### Immunohistochemistry analysis

All slides were digitalized using the Aperio Scanscope XT slide scanner. Evaluation of stained tissue slides was performed using QuPath (version 0.2.0) software. The percentage of tumor tissue binding NC410 was calculated by annotation of the tumor in the tissue slide and quantifying the percentage of DAB-stained tissue by using the Pixel Classifier.

Scoring of the immune phenotype was based on the presence and distribution of CD3 positive lymphocytes(61). Tumors with CD3^+^ cells in the tumor fields were scored “inflamed”; tumors with CD3^+^ cells in their stroma, but without or with a relatively low amount of CD3^+^ in the tumor fields were scored “immune excluded”; tumors with a lack of CD3^+^ cells in their stroma as well as in their tumor fields were scored “immune desert”.

For quantification of the immune cell counts, regions of interest (ROIs) were annotated on the tissue slides by drawing circles with a diameter of 600 μm at five random spots within the NC410 binding part of the tumor. Positive cells were quantified using the Positive Cell Detection tool. For each staining, stain vectors and DAB cutoffs were determined based on a representative slide; the settings were kept the same for all slides. Tumor area within the circles that was lost during staining procedure or that was negative for NC410 binding was excluded from the analysis.

The percentage of NC410 binding to tissue within a tumor was calculated by dividing the stained area by the total tumor area. The immune cell counts for each tumor were calculated by dividing the total number of positive cells within the five ROIs by the total surface in mm^2^ of these ROIs.

### Statistics

Data were graphed and analyzed using GraphPad Prism 8.0 (Graph Pad software). Data are presented as the mean ± standard error of the mean (SEM). The significance was analyzed with unpaired, two-tailed Student’s t-tests and 2-way ANOVA followed by multiple comparison.

## Acknowledgments

We thank Michiel van der Vlist for his critical comments on the manuscript.

## Competing interests

LT, CS, AP, JS, JB, ZC, LL, SL and DF are employees from Nextcure. Nextcure holds a patent on NC410.

## Author Contributions

**Conceptualization:** ZC, LL, SL, SW, DF, LM

**Data curation:** JS

**Formal analysis:** MIPR, LT, EJR, CS, AP, AS, EE, SVV, JS, LL, SL, SW, DF, LM

**Funding acquisition:** DF, LM

**Investigation:** MIPR, LT, EJR, CS, AP, AS, EE, SVV, JS, JB

**Methodology:** MIPR, LT

**Project administration:** ZC

**Supervision:** DF, LM

**Writing – original draft:** MIPR, LT

**Writing – review & editing:** MIPR, LT, EJR, CS, AP, AS, EE, SVV, JS, JB, ZC, LL, SL, SW, DF, LM

---

**Supplemental Figure 1.**
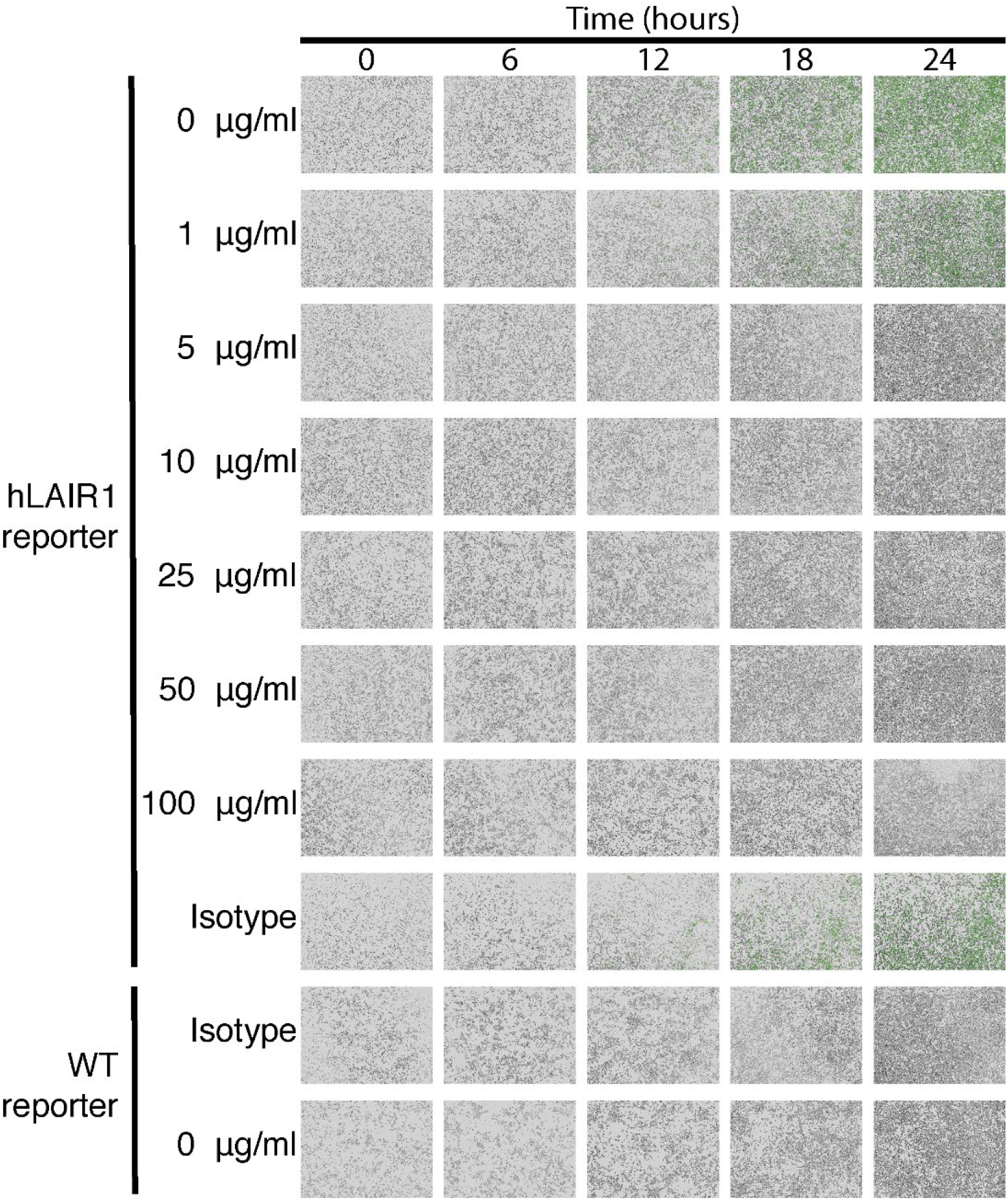
NC410 blocks LAIR-1 functional interaction with collagen. NC410 dose response measured by incucyte imaging during 24h. Representative microscopic images (10x) of WT and hLAIR-1 reporter cells over time. NC410 protein was added at increasing concentrations and inhibited collagen I (5 ug/mL) mediated NFAT-GFP signaling through LAIR-1 binding. Isotype control was used at the highest concentration (100 ug/mL) and showed no inhibition of NFAT-GFP signaling. Pictures were taken every 1 h for 24 h. Representative images from 1 out of 3 independently performed experiments (each with experimental triplicates).

**Supplemental Figure 2.**
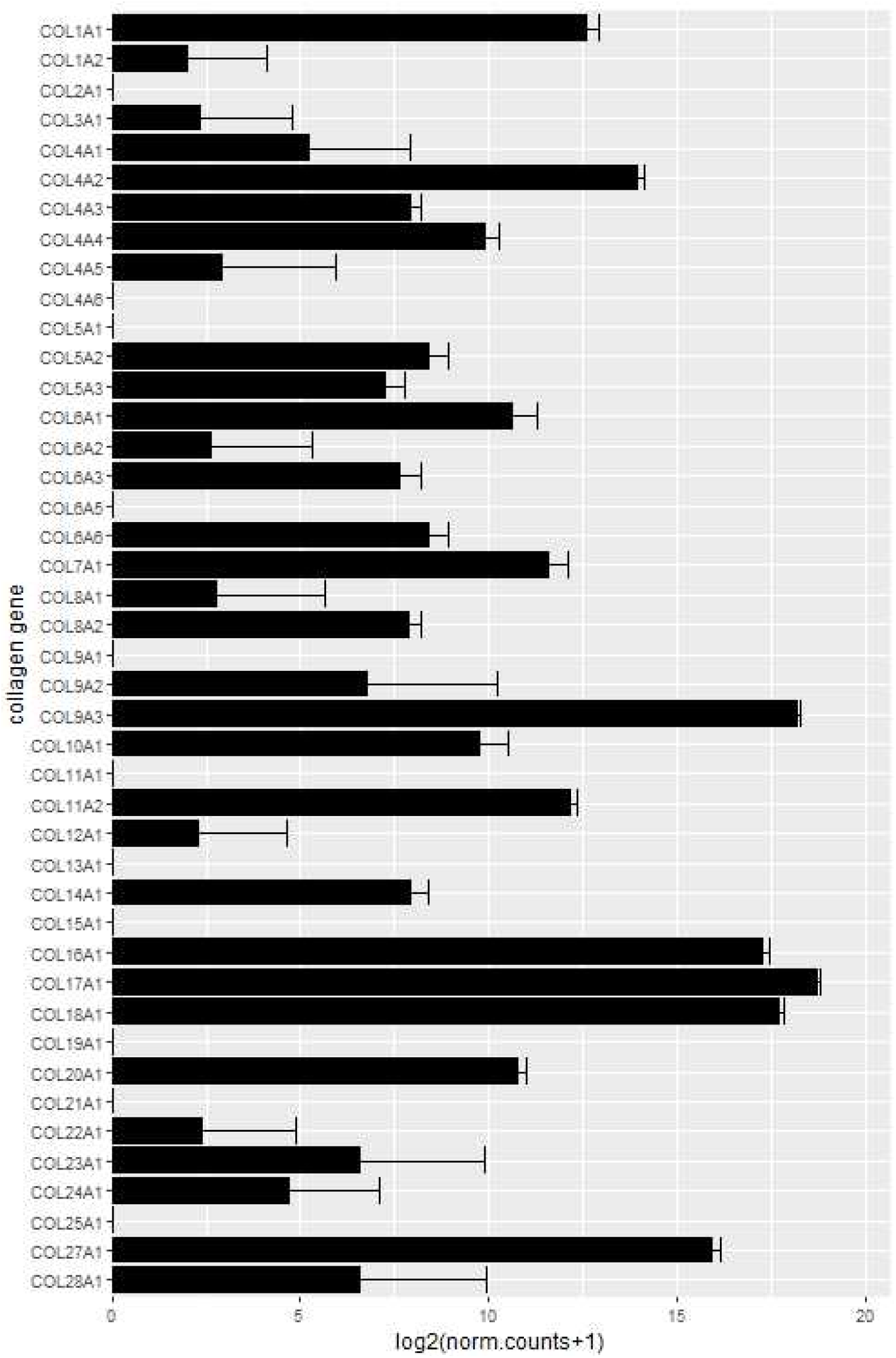
HT-29 mRNA collagen expression by RNA sequencing. For collagen gene expression analysis of HT-29 cells, data was acquired from the public dataset GSE41586 (https://pubmed.ncbi.nlm.nih.gov/23902433/). Raw count data of untreated HT-29 cells was retrieved and normalized using the DESeq2 package (v1.28.1) in R (v4.0.2). Data was then log2 transformed and plotted using the ggplot package (v3.3.2).

**Supplemental Figure 3.**
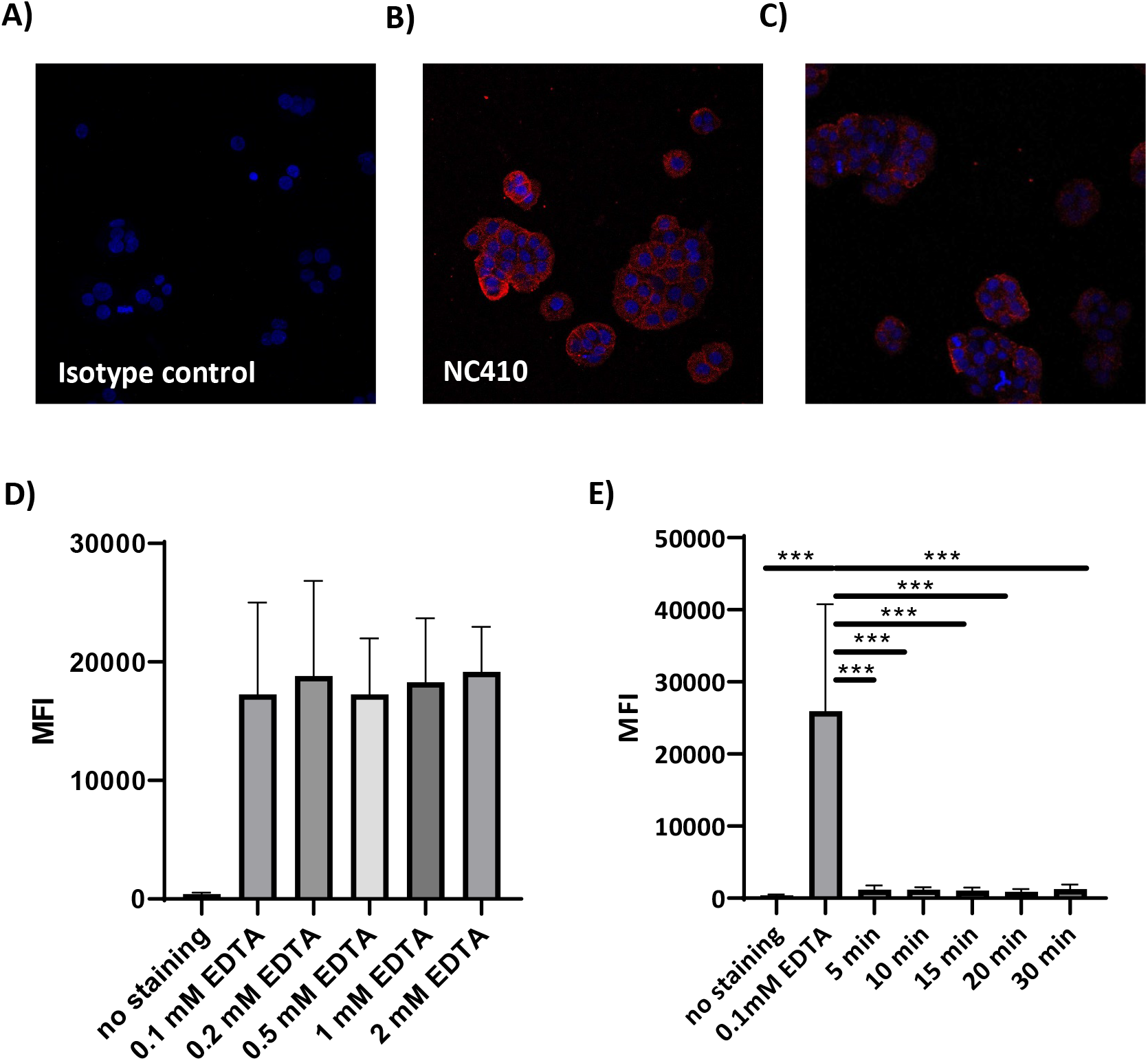
HT-29 collagen expression. Immunofluorescence analysis of HT-29 cells stained with A) isotype control, B) NC410 or C) pan-collagen antibody. 40x magnification D) HT-29 cells were removed from culture flasks using increasing concentrations of EDTA for 10 min. MFI of NC410 staining is shown, demonstrating that EDTA treated HT-29 cells keep surface collagen expression. E) HT-29 cells treated with 0.1mM collagenase lose NC410 binding. Data from 3 independently performed experiments.

**Supplemental Figure 4.**
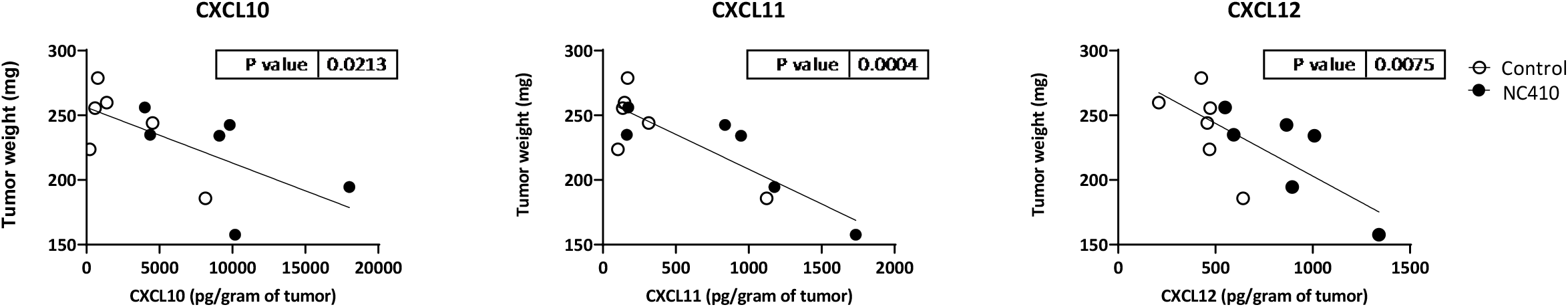
Chemokine expression in a humanized HT-29 tumor model. CXCL10, CXCL11 and CXCL12 chemokine expression correlate with reduced tumor growth at day 27 after HT-29 injection. P value was calculated by linear regression with F test. 6 mice per group were used. Black circles indicate NC410 treatment and open circles indicates control treatment.

**Supplemental Figure 5.**
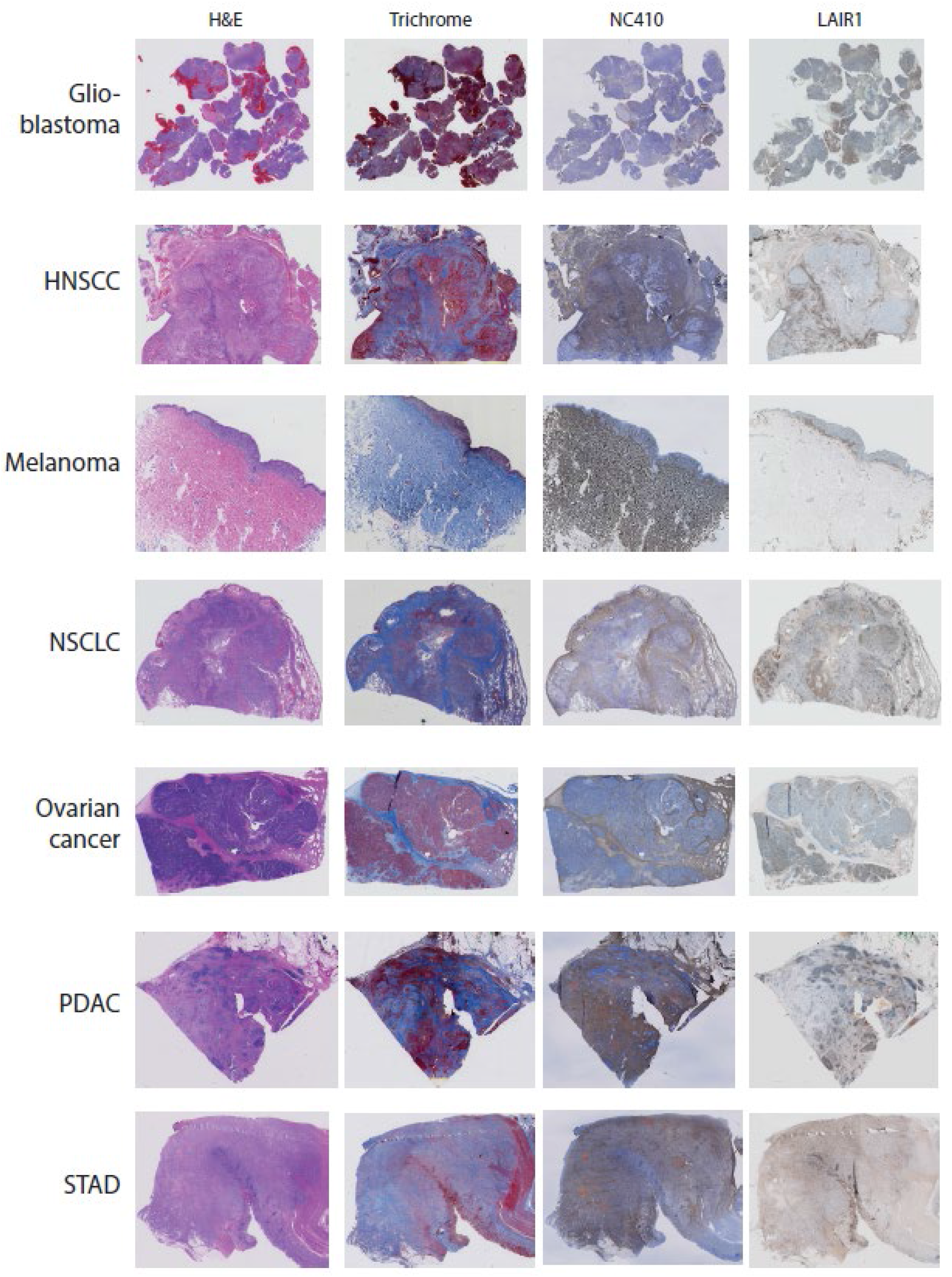
Immunohistochemical analysis of primary human tumors. Representative hematoxylin and eosin (H&E), Masson Trichrome, NC410 and LAIR-1 staining for 7 different tumor types (Head and neck (HNSC), melanoma, -non-small cell lung carcinoma (NSCLC), ovarian, pancreatic (PDAC) and stomach cancer (STAD))

**Supplemental Figure 6.**
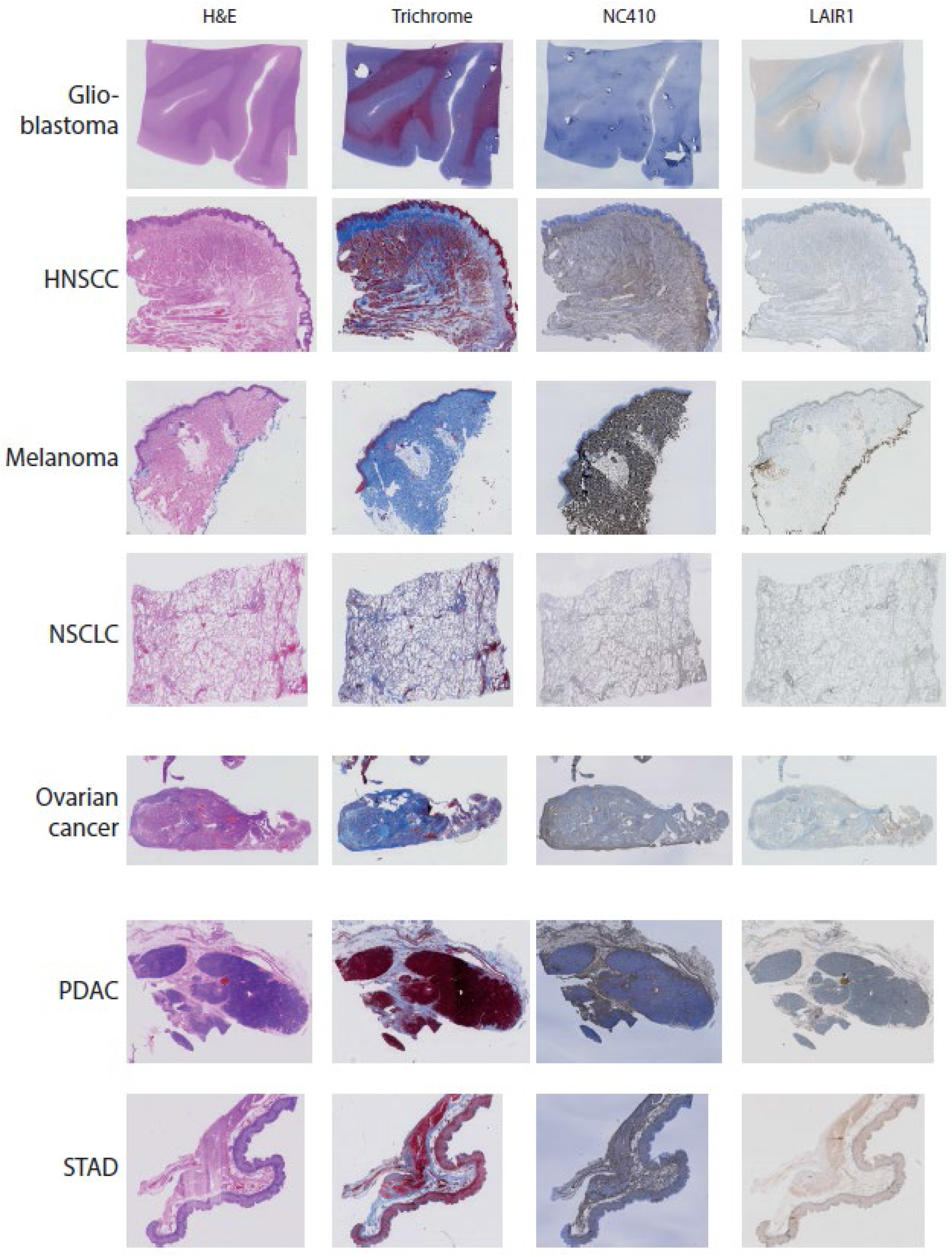
Immunohistochemical analysis of healthy tissue. Representative hematoxylin and eosin (H&E), Masson Trichrome, NC410 and LAIR-1 staining for healthy tissue matching the tumors used in Supplemental Figure 5.

**Supplemental Figure 7.**
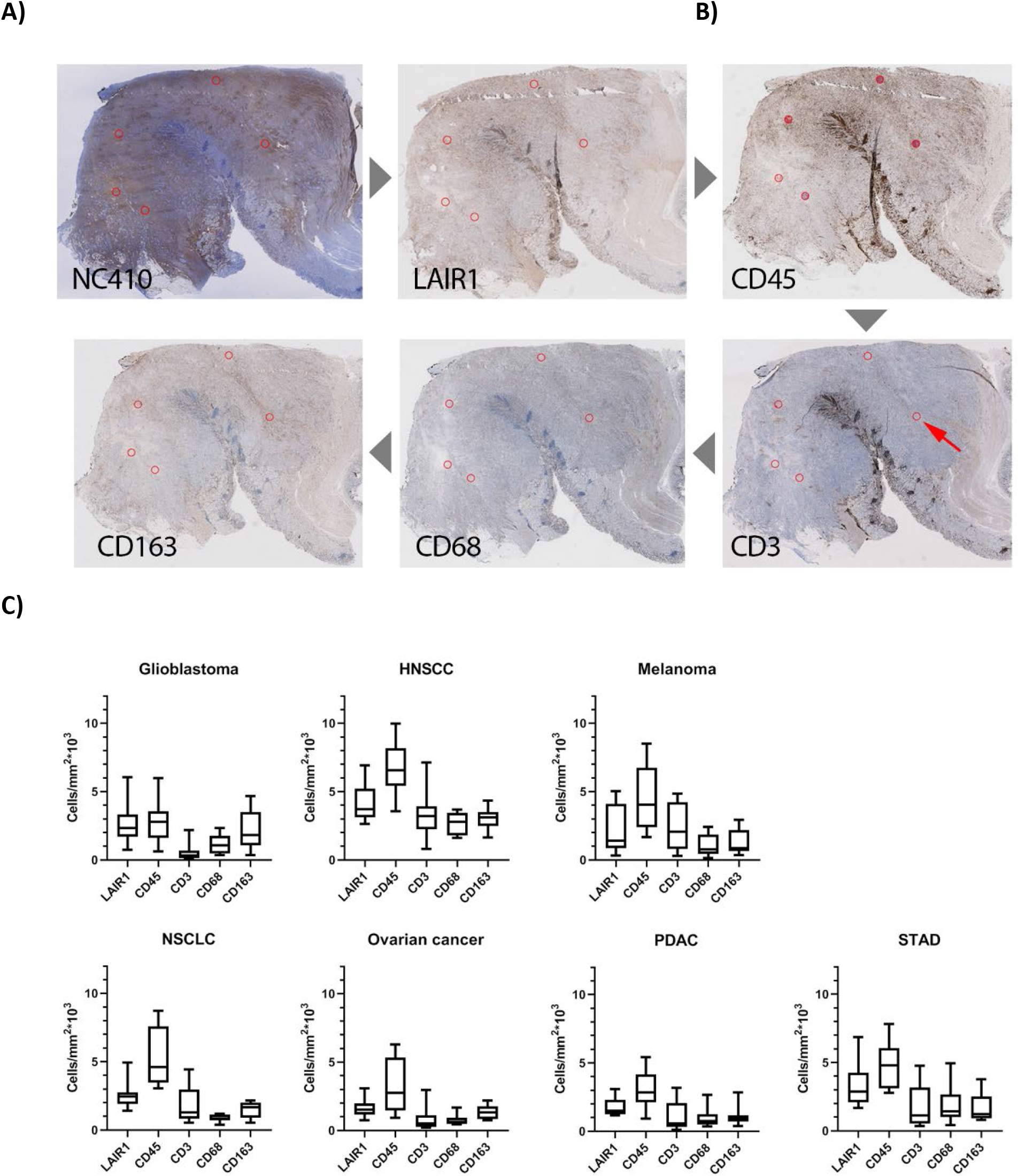
Characterization of LAIR-1^+^ cells in primary human tumors. A) Representative LAIR-1 FC, NC410, CD45, CD3, CD68 and CD163 staining in a stomach tumor specimen. B) For quantification of the immune cell counts, regions of interest (ROIs) were annotated on the tissue slides by drawing circles with a diameter of 600μm at five random spots within the NC410 positive part of the tumor. Positive cells were quantified using the Positive Cell Detection tool. C) Number of positive LAIR-1 FC, CD45, CD3, CD68 and CD163 cells are shown across 7 different tumor types (Head and neck (HNSC), melanoma, -non-small cell lung carcinoma (NSCLC), ovarian, pancreatic (PDAC) and stomach cancer (STAD)).

**Supplemental Figure 8.**
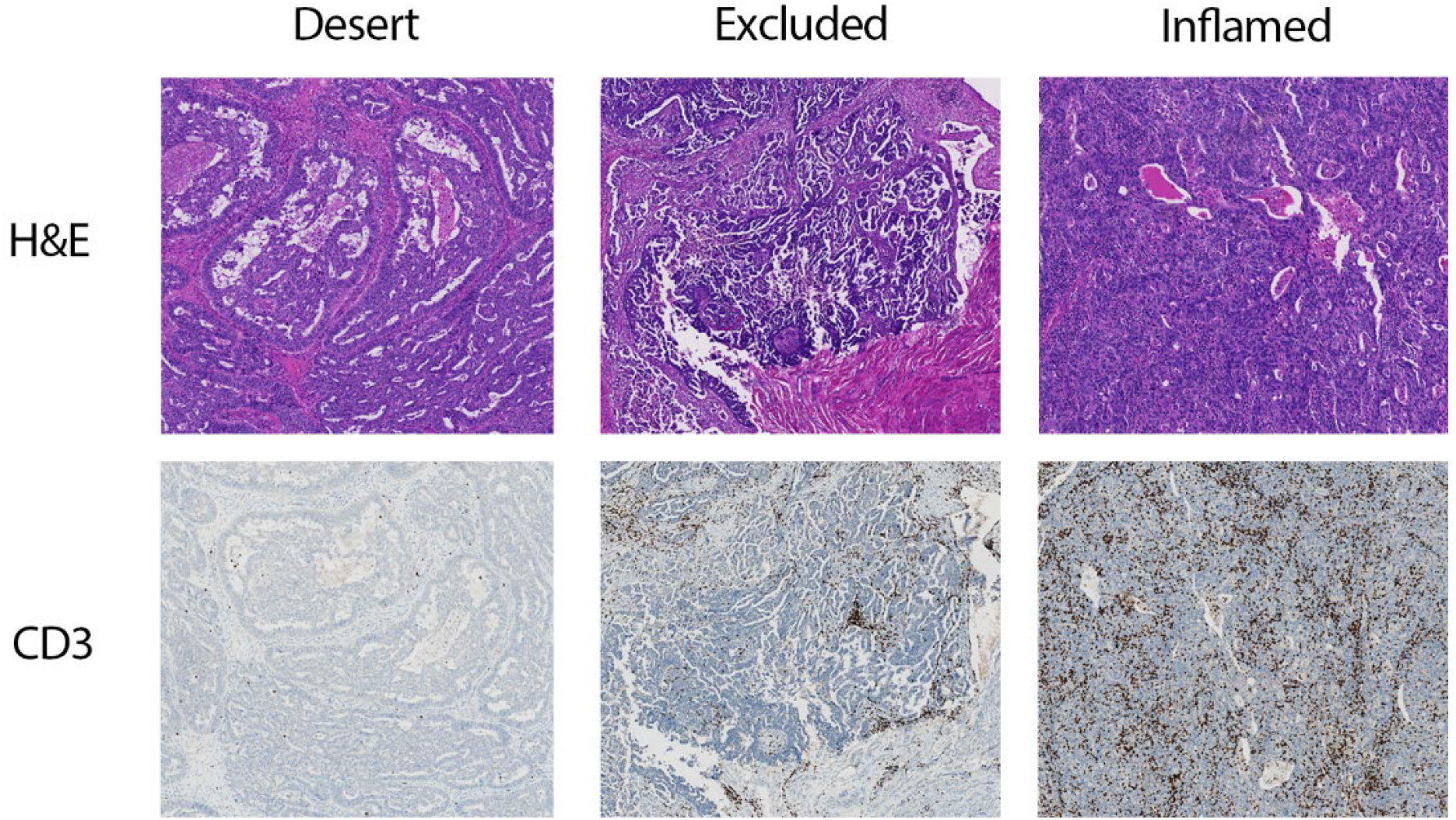
Tumor immune phenotyping. Representative image of the 3 different immune phenotypes, immune desert, immune excluded and immune inflamed in an ovarian tumor specimen.

